# Vertical transmission of mosquito microbiota and its effects on offspring development

**DOI:** 10.64898/2025.12.16.694692

**Authors:** Holly L. Nichols, Laura E. Brettell, Miguel A. Saldaña, Shivanand Hegde, Eva Heinz, Kathy Vaccaro, Grant L. Hughes, Kerri L. Coon

## Abstract

**Background:** The mosquito’s gut microbiota plays a crucial role in determining its capacity to transmit harmful viruses and parasites. Accordingly, manipulating mosquito gut microbiota is a promising avenue towards reducing mosquito-borne human pathogen transmission. A successful microbial control campaign will require a thorough understanding of how bacteria are transmitted through mosquito populations. Through two parallel but complementary studies using the yellow fever mosquito *Aedes aegypti* as a focal host species, we surveyed vertical transmission of bacteria from individual mothers to cohorts of offspring maintained in a closed system.

**Results:** Laboratory- and field-derived mothers deposited bacteria that support offspring development, and the relative abundance of commonly transmitted taxa correlated with offspring fitness. Maternally transmitted bacteria were detected in both larval and adult offspring, and the relative abundances of specific taxa differed between life stages. Microbiota composition in adult offspring closely resembled microbiota composition in mothers, despite dramatic shifts in the relative abundance of specific microbial community members during the larval stage. Variability in microbiota composition in offspring was also greater than variability across the population of egg-laying mothers. Eggs that underwent a period of desiccation before hatching produced larval communities dominated by endospore-forming bacteria that were rare in maternal samples.

**Conclusions:** Overall, our results demonstrate the vertical transmission of mosquito-associated microbiota across generations, including bacterial taxa that could potentially be leveraged for mosquito and mosquito-borne disease control.

## Background

Mosquitoes are the primary vectors of numerous human pathogens, and morbidity and mortality from mosquito-borne diseases are predicted to increase due to land use practices and climate change [1, 2]. Classic vector control practices such as the use of broad-spectrum insecticides have reduced disease burden, but growing insecticide resistance and continued disease transmission even in well-managed districts suggests that current techniques are insufficient to surmount the growing need for vector control [3]. New technologies are required as part of an integrated management campaign to stop the spread of mosquito-borne pathogens.

Mosquitoes have an internal gut microbial community, or gut microbiota, that makes vital contributions to the physiology of the mosquito host. The gut microbiota are implicated in traits directly associated with pathogen transmission such as vector competence and insecticide resistance [4–6]. Gut microbes also contribute to fundamental aspects of mosquito physiology; live microbes are required for development past the first instar (L1) under natural conditions due to their vital contribution of metabolites such as riboflavin [7, 8]. Community assembly has downstream effects on mosquito development time, nutrient stores, blood meal digestion and fecundity, all of which contribute to vector population dynamics [9]. Consequently, there is profound interest in understanding how gut bacterial communities are acquired and how they persist throughout the mosquito life cycle.

Mosquitoes begin their life cycle as aquatic larvae, pupate in larval water, and emerge as terrestrial adults that seek out standing water to lay eggs [10]. Larvae hatch with a sterile digestive tract and acquire gut microbes via browsing on egg casings and feeding in their aquatic habitat [11]. As such, larval microbial communities closely resemble the microbial community of the surrounding water, although gut community diversity is generally lower [12–17]. Larval microbial communities subsequently experience frequent turnover as larvae grow and molt through four instars before pupating on the water’s surface. When larvae pupate, gut contents are egested, and the diversity and abundance of any remaining gut microbiota are further reduced as gut tissues are remodeled during metamorphosis to the adult stage. Adult mosquitoes therefore emerge with relatively depauperate gut microbial communities as compared to larvae [18–20]. The gut is then quickly recolonized as individuals of both sexes imbibe larval water and forage for nectar in the first days after emergence [21, 22]. Females of vector species undergo an additional perturbation when they take a blood meal, which causes an increase in bacterial load and a decrease in diversity when bacteria unable to tolerate the oxidative stress are excluded [23, 24]. These general patterns of acquisition and community assembly have been well-studied in various host species in both laboratory and field populations [25]. The fidelity of microbiota between generations, however, is less clear.

Microbial community members are maintained in host populations by a combination of horizontal and vertical transmission between individuals. In insects, microbes are passed vertically between generations in a variety of ways, including inside the egg, on the egg surface, or by inoculating the environment where offspring develop [26, 27]. Egg-smearing, or vertical transmission of bacteria via deposition on egg surfaces, has been observed by allowing mosquitoes to lay eggs under sterile conditions, but it remains unclear which proportion of the material bacterial community persists in subsequent generations [7, 27, 28]. Although vertical transmission of bacteria could be hindered by diverse bacterial consortia naturally occurring in breeding water, two lines of evidence suggest that maternally deposited microbes persist in subsequent generations. First, studies following individual fluorescently-labelled isolates demonstrate transmission from adult females to egg surfaces and even persistence through adulthood for *Serratia*, *Asaia* and *Pantoea* [11, 29, 30]. Second, a recent study found that egg laying (oviposition) alters larval water microbial communities in a manner distinct from simple physical contact between female mosquitoes and the water, implying indirect vertical transmission via inoculation of larval water [31]. For these reasons, we posit that maternal transmission of microbiota is an important factor shaping mosquito microbial communities and fitness outcomes.

Here, we completed two independent but complementary experiments to characterize the contribution of vertical (maternal) transmission to offspring development and microbiota assembly, using the yellow fever mosquito *Aedes aegypti* as a focal host species. *Ae. aegypti* is an ideal model system for such experiments due to its role as a primary vector of arboviruses causing disease in humans. We have also recently developed tools for microbiota isolation, cryopreservation, and transplantation in this species [23, 32–38], which allowed us to conduct our experiments in a closed system and isolate the effects of microbial inputs from individual ovipositing females. In the first experiment, we tracked time to pupation, survival to adulthood, and wing length for offspring immediately hatched from eggs laid by individual laboratory-reared female mosquitoes and correlated those outcomes with abundant bacterial taxa. In the second experiment, we included ovipositing females from both laboratory and field populations and tracked microbial community composition through adulthood for offspring that hatched from eggs stored for three months. Our results have important implications for our understanding of (i) the bacterial assemblages that are vertically transmitted from ovipositing adult females to offspring in *Ae. aegypti* mosquitoes, (ii) the degree of variation in material transmission of mosquito microbiota within a population, (iii) the effect of egg storage on vertically transmitted bacterial communities, and (iv) the fitness impacts of maternally transmitted microbes. This information could be leveraged to improve ongoing efforts to identify candidates for microbe-based mosquito and mosquito-borne disease control, as control campaigns utilizing microbes that persist across host generations will be effective beyond the lifespan of targeted individuals.

## Methods

### Mosquito rearing and experimental setup – Experiment 1

Experiment 1 was conducted at the University of Wisconsin-Madison (UW-Madison; Fig 1a). In brief, the Liverpool (LVP) strain of *Ae. aegypti* was conventionally reared in an established insectary under the following environmental conditions: constant temperature of 26.5 ± 1 °C, 75 ± 10% relative humidity, and a 16:8 hr light: dark photoperiod with 90 min crepuscular periods. Larvae were hatched in deoxygenated water. Three replicate pans (A, B, and C) of L1 larvae were reared at a density of 300 larvae per aluminum rearing pan and kept in distilled water and fed ground tropical fish flakes (Tetra, Melle, Germany) *ad libitum*. Pupae were collected and allowed to emerge in non-sterile 473 mL/16 oz cartons (Solo Cup Company, Lake Forest, IL USA) at a ratio of 35 females to 17 males. Upon emergence, adults were provided with 5% (wt/vol) sterile sucrose solution *ad libitum*. Females received defibrinated sheep blood (Hemostat Laboratories, Dixon, CA USA) on Day 4 post-emergence.

**Figure 1:**
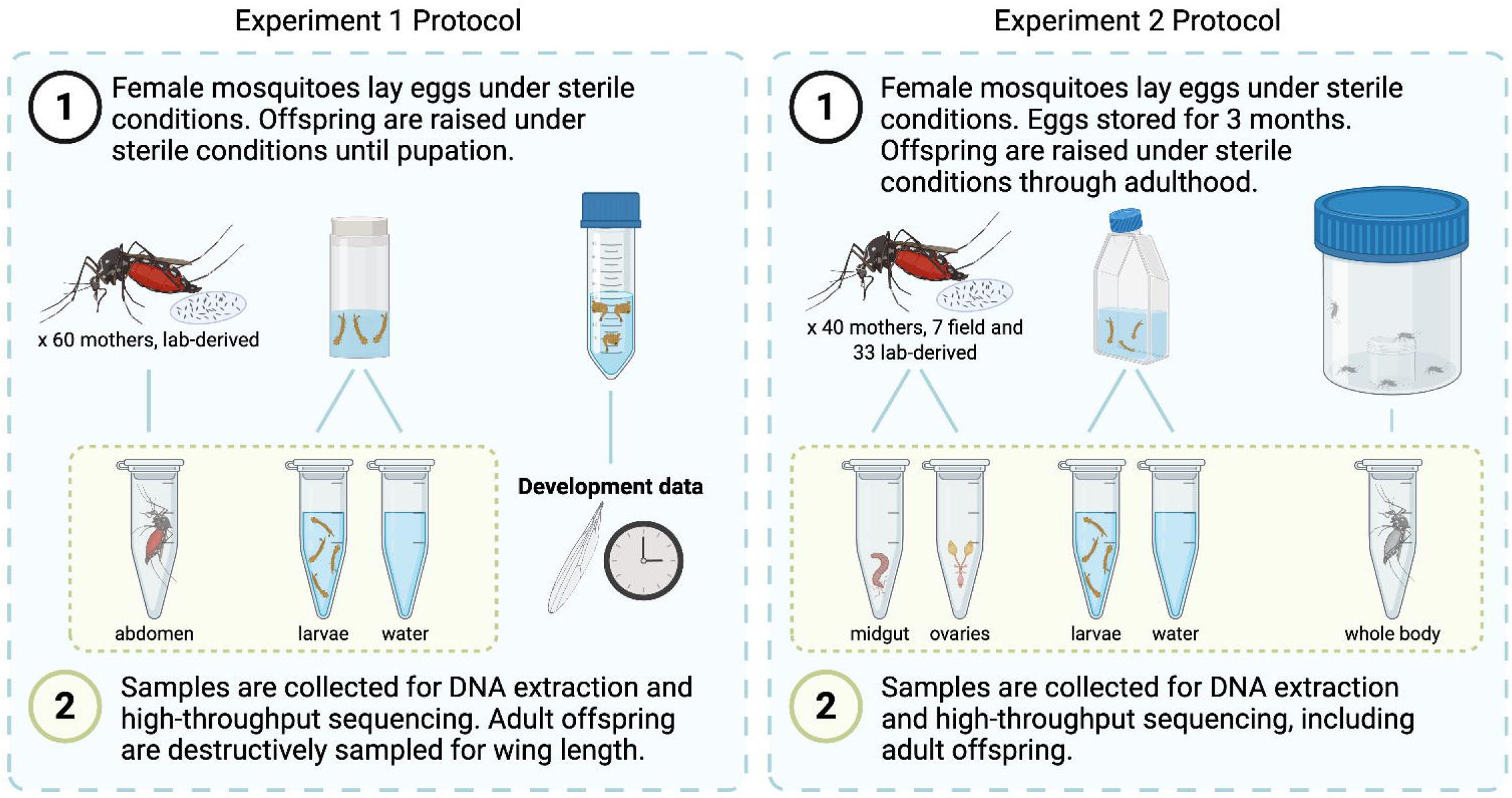
Comparison of methodology between Experiment 1, which was conducted at the University of Wisconsin-Madison (Madison, WI, USA), and Experiment 2, which was conducted at the University of Texas Medical Branch (Galveston, TX, USA). In both experiments, non-sterile *Ae. aegypti* (Liverpool, LVP, or Galveston, GALV, strain, respectively) adult females laid eggs under sterile conditions. Eggs in Experiment 1 were immediately hatched, while eggs in Experiment 2 were stored for 3 months at ambient laboratory conditions. Offspring were then reared under sterile conditions until an endpoint. In Experiment 1, whole abdomens of ovipositing females were collected and processed for downstream 16S rRNA gene amplicon sequencing. In Experiment 2, ovary and midgut tissues were dissected from each ovipositing female and processed separately for sequencing. Larval offspring and an aliquot of larval water were also sampled and processed for sequencing in both experiments. While adult female offspring in Experiment 1 were sacrificed to measure wing length as a proxy for adult quality, adult female offspring in Experiment 2 were sampled at 11 days post-emergence and processed (without dissection) for sequencing. Overall, Experiment 1 included 60 laboratory-reared *Ae. aegypti* LVP females, from which 28 isofemale lines were taken through to library preparation and sequencing; Experiment 2 included 7 field-caught *Ae. aegypti* and 33 lab-reared *Ae. aegypti* GALV females, of which 3 and 7 isofemale lines, respectively, were taken through to sequencing. Figure created with BioRender.com.

Approximately 72 hr post-blood meal, all mosquitoes were cold anesthetized at 4 °C for 10 min and individual females were moved in a biosafety cabinet to sterile oviposition chambers containing sterile filter paper wet with sterile water. Sixty F0 females (hereafter referred to as “mothers”) were allowed to oviposit in these chambers for 24 hr at 28 °C with a 16:8 hr light: dark photoperiod. After 24 hr, mothers were stored in individual microcentrifuge tubes at −80 °C for further processing. Egg papers were immediately floated in culture flasks containing 30 mL of sterile water and 2.75 mg of gamma-irradiated sterile rat chow, lactalbumin and dead torula yeast (1:1:1), ensuring any input of microbes on the eggs was deposited by the female. Eggs were incubated for 72 hr, and then a vacuum was applied for 45 min to synchronize hatching. Newly hatched F1 larvae were kept in culture flasks for an additional 24 hr to ensure colonization by microbes in the water. In parallel, three lines were additionally hatched from egg sheets collected from the insectary population to represent potential microbial inputs at a population level. These egg sheets had been stored in Whirl-Pak bags (Nasco, Fort Atkinson, WI) for approximately three months under aforementioned insectary conditions before hatching, as is standard practice for colony maintenance. These “insectary control” lines were maintained identically to the isofemale lines after hatching.

Approximately 24 hr post-egg hatching, larvae were rinsed three times with sterile water and approximately 20 individual larval offspring per mother were moved to an autoclaved *Drosophila* vial (VWR International, Radnor, PA USA) containing 10 mL sterile water. Flasks that did not yield at least 20 larvae were discarded, resulting in 29 isofemale lines. An additional three tubes were maintained without larvae and were fed and sampled identically to experimental tubes; water from these negative control tubes was plated on LB (Miller) agar every three days to confirm the absence of culturable contaminants. Larvae were fed a diet of sterile rat chow mix, according to a previously optimized feeding regimen to support development to the adult stage [39]. F1 larval water and larval offspring were sampled and frozen at −80 °C at the following time points throughout the experiment: (*i*) 10 mL larval water 24 hr post-egg hatching, (*ii*) pools of 5-10 first instar (L1) larvae 24 hr post-egg hatching, (*iii*) 1 mL larval water 72 hr post-egg hatching, (*iv*) one fourth instar (L4) larva on the day of first pupation in each tube, (*v*) 1 mL larval water on the day of first pupation in each tube, and (*vi*) 1 mL larval water eighteen days after hatching (Additional File 1). All pupae resulting from a given cohort of larvae were rinsed once with sterile water and moved in 20 mL sterile water to a sterile 50 mL conical tube (Thermo Fisher Scientific, Waltham, MA USA) for F1 adult emergence. Adult offspring were removed from conical tubes shortly after emergence and the size of each individual was determined by measuring the length of one forewing using a stage micrometer as described below.

### Mosquito rearing and experimental setup – Experiment 2

Experiment 2 was conducted at The University of Texas Medical Branch (UTMB; Fig 1b), with analyses conducted at the Liverpool School of Tropical Medicine (LSTM). Here, both laboratory-reared and field-caught *Ae. aegypti* were used to generate isofemale lines. For the laboratory-reared mosquitoes, Galveston (GALV) line *Ae. aegypti* were used, which had been continually maintained at the UTMB at 28 °C with a 12:12 hr light: dark photoperiod. Larvae were fed finely ground pellets of fish food (Tetramin), and adults were provided with 10% (wt/vol) sucrose solution *ad libitum.* In brief, thirty-three individual blood-fed F0 females (hereafter referred to as “mothers”) from the same batch of eggs were transferred to sterile *Drosophila* vials containing strips of filter paper wet with sterile water for oviposition. Females were blood fed 3-5 days post eclosion on sheep’s blood (Colorado Serum Co.) until fully engorged. For the field-caught mosquitoes, collections were made locally in Galveston, Texas using Biogents BG-sentinel 2 traps in October of 2017. Adult mosquitoes were collected and sorted morphologically according to species and sex. Seven F0 field-caught female *Ae. aegypti* mothers were selected and transferred individually to separate sterile containers for oviposition as described above.

Six days post-blood feeding, vials containing mothers that had not oviposited were discarded. Mothers that had successfully laid eggs were then removed from vials and anesthetized by storage at 4 °C for 5-10 min. These F0 mothers were then surface sterilized by a series of sequential washes (5 min in 70% ethanol, 5 min in 10% bleach, three 5-min washes in sterile phosphate buffered saline solution) and their guts and ovaries were dissected with sterile forceps before storing at −80 °C. Three replicate samples of oviposition water were also collected and stored at −80 °C. Egg papers were collected and stored for three months in sterile bags in accordance with standard insectary colony maintenance techniques. The number of eggs laid per adult was recorded and egg papers were then transferred to sterile T175 culture flasks (Fisher) with 140 mL sterile water and autoclaved finely ground fish food (Tetramin). F1 larval offspring were allowed to hatch and develop until most individuals had reached the 3^rd^/4^th^ instar (L3/L4) (or until 20 days post-egg hatching). At this point, ten L3/L4 F1 larvae were surface sterilized and stored individually at −80 °C (Additional File 1). Three replicate 200 μL samples of F1 larval water and three replicates of fish food slurry were also collected and stored at −80 °C. The remainder of the larvae were allowed to continue development and at the pupal stage were transferred to sterile tissue culture roller bottles. Emerging F1 adult offspring were provided 10% (wt/vol) sterile sucrose solution *ad libitum* and, at 11 days post-transfer, adults were counted, and all females were collected and stored individually at −80 °C. As outlined above, multiple isofemale lines of laboratory-reared (*n* = 33) and field-caught mosquitoes (*n* = 7) were assayed. However, only 10 of these lines (seven from laboratory-reared and three from field-caught mosquitoes) yielded sufficient offspring for sequencing at each time point. Successful lines were selected 7 days post-egg hatching.

### Sample processing, DNA extraction, and 16S rRNA gene amplicon sequencing – Experiment 1

For Experiment 1, total genomic DNA was extracted from all frozen samples using phenol-chloroform as described previously [38, 40, 41]. Prior to DNA extraction, water samples were subjected to two minutes of mechanical disruption in 800 µL of extraction buffer using 425-600 µM acid-washed and autoclaved glass beads (BioSpec Products, Bartlesville, OK USA) and a bead beating machine. Mother samples were rinsed for 10 sec in 70% ethanol followed by three rinses in sterile water, and whole abdomens were dissected. Larval offspring samples were rinsed three times with sterile water. Both mother and larval offspring samples were then homogenized with a single 5-mm carbon-steel bead (Qiagen, Hilden, Germany) for 20 sec to disrupt cuticle tissues and subjected to two minutes of bead-beating with glass beads as described above. Extraction blanks were generated using nuclease-free water as template (*n* = 7).

Approximately 10 ng of DNA from each sample was subjected to one-step PCR amplification of the V4 region of the bacterial 16S rRNA gene using 2x HotStart Ready Mix (KAPA Biosystems, Wilmington, MA USA) and barcoded primers as described previously [35, 39]. Reactions containing DNA from F0 mother samples were amplified at 35 cycles, while reactions containing DNA from F1 larval water and larval offspring samples were amplified at 25 cycles. No-template reactions and reactions using template from blank DNA extractions served as negative controls. PCR products were visualized on 1% agarose gels and successful amplifications were purified using a MagJET NGS Cleanup and Size Selection kit (Thermo Fisher Scientific, Waltham, MA USA). DNA concentration in the resulting 253 purified libraries was finally measured using a Quantas fluorometer (Promega, Madison, WI, USA) and an equimolar pool of all libraries was submitted for paired-end (2 x 250 bp) sequencing on an Illumina MiSeq by the DNA Sequencing Facility at the UW-Madison (Madison, WI USA).

### Sample processing, DNA extraction, and 16S rRNA gene amplicon sequencing – Experiment 2

For Experiment 2, total genomic DNA was extracted from different sample groups using a NucleoSpin Tissue kit (Machery-Nagel, Düren, Germany) as follows: F1 larval water, oviposition water, and fish food slurry samples were first mixed with 180 μL T1 lysis buffer plus 20 μL 100 mg/mL lysozyme and incubated at 37 °C for 1 hr; 25 μL proteinase K was then added and the samples were incubated at 56 °C for an additional 3 hr. DNA extractions continued following manufacturer instructions. F0 mother midgut and ovary samples were first homogenized for 5 min at 60 Hz/sec in a TissueLyzer (Qiagen) in 80 μL T1 buffer plus 10 μL 100 mg/mL lysozyme and incubated at 37 °C for 1 hr; 8 μL proteinase K was then added and the samples were incubated at 56 °C overnight. DNA extractions then continued following manufacturer instructions, with the following volume modifications: 80 μL B3 buffer, 80 μL 100% ethanol, 50 μL B5 buffer, and elution in 20 μL nuclease-free water at 70 °C. Surface-sterilized F1 adult female offspring were first homogenized in 180 μL T1 lysis buffer with 25 μL proteinase K and a single sterile stainless-steel bead (Qiagen) using a TissueLyser (Qiagen, Hilden, Germany) set to 30 Hz for 30 sec. Homogenates were then incubated at 56 °C overnight before proceeding according to manufacturer instructions.

With each batch of extractions, blanks were also included, using nuclease-free water (*n* = 5). DNA was quantified by spectrophotometry (Nanodrop) and shipped on dry ice to the Centre for Genome Innovation at the University of Connecticut (Mansfield, CT USA) for library preparation using primers targeting the V3-V4 region of the bacterial 16S rRNA gene (515F and 806R) and sequencing on an Illumina HiSeq to generate 250-bp paired-end reads.

### Sequencing data analyses – Experiments 1 & 2

Sequencing analyses were completed in tandem by scientists at the UW-Madison and the LSTM. For the data generated in Experiment 1, de-multiplexed reads were imported into QIIME for quality control and taxonomic assignment as described previously [38, 42]. In brief, DADA2 was used to denoise reads [43]. Taxonomy was assigned with a Naïve-Bayes classifier derived from the Silva-138-99 database (accessed June 16, 2022) [44]. Multiple sequence alignment and phylogenetic tree construction were performed using mafft and FastTree2, respectively [45, 46]. Endpoint .qza files from QIIME2 were imported into R version 4.1.3 (https://www.r-project.org) and merged with metadata into a phyloseq object for further processing [47]. Reads identified as Chloroplast, Mitochondria, or any reads without a domain identification of “Bacteria” were discarded. Detection and elimination of contaminant sequences was carried out using the “prevalence” and “frequency” methods in the *decontam* package [48]. After manual inspection of putative contaminants, one amplicon sequencing variant (ASV), *Elizabethkingia*_b61, was retained due to its abundance (>95% of reads) in 11 true samples and rarity in control samples. Samples with fewer than 100 sequences after decontamination were eliminated from further processing, which eliminated six out of 28 mother samples. For some analyses, samples of each type (mother, water, larvae) from the same isofemale line were merged. Merged data was then passed back to QIIME2, where alpha diversity (measured by Shannon’s *H* index) and beta diversity (measured by the unweighted Unifrac and Bray-Curtis distance metrics) were calculated using standard workflows and a p-sampling-depth of 8,599 according to the plateauing of the rarefaction curve generated using the ‘ggrare’ function in the *ranacapa* package in R [49] (Additional file 2), which excluded one sample from alpha and beta diversity analysis. Pertinent .qza files were exported to R for visualization and statistical analysis.

For the data generated in Experiment 2, the same method was used for sample QC and filtering, with the following exceptions: after using the “prevalence” method in *decontam* to identify and remove potential contaminants (using a threshold of 0.5 following recommendation of Diaz *et al.* [50]). To ensure that life-stage-specific taxa were not erroneously removed, ASVs were retained if their average relative abundance any sample type (ovaries, midguts, larval water, larval offspring, adult offspring) exceeded that in control samples. For alpha and beta diversity analysis, samples were then rarefied to a depth of 1,005 reads following inspection of rarefaction curves (Additional file 3), with any samples containing fewer reads being discarded. In Experiment 2, data generated from larval and adult offspring were analyzed at the individual level. Again, alpha and beta diversity metrics were calculated in QIIME2 on the rarefied data, with statistical analyses and visualization conducted in Rstudio v2022.07.1.

For both datasets, differences between the alpha diversity (Shannon’s *H* Index) were assessed via Kruskal-Wallis rank-sum tests using the ‘kruskal.test’ function in the *vegan* R package [51], with *post-hoc* Bonferroni-corrected pairwise comparisons calculated via a Dunn’s test implemented with the *dunn.test* package [52]. Differential dispersion of samples by type was analyzed using PERMDISP2 followed by a pairwise permutation-based test of multivariate homogeneity of group dispersions (permutest, pairwise.permutest). Clustering by sample type was analyzed by permutational multivariate analysis of variance (PERMANOVA) with the ‘adonis2’ function in the *vegan* package [51], followed by *post-hoc* comparisons using the ‘pairwise.adonis2’ function in the *pairwiseAdonis* package [53]. Differential abundance analysis was conducted at the ASV level with the ALDEx2 package; t.test and effect modules were employed for Wilcoxon rank sum tests between two groups and the aldex.kw module was used to conduct Kruskal-Wallis tests for three or more groups [54]. Custom R scripts were generated for tracking ASVs between paired mother and offspring samples and are available on GitHub (https://github.com/kcoonlab/mosquito-vertical-transmission).

### F1 fitness measurements and data analysis – Experiment 1 & 2

F1 development outcomes from Experiment 1 were quantified via two parameters: duration of larval development (days from egg-hatching to pupation) and adult body size (approximated by wing length). Wings of emerged F1 adult offspring were mounted on glass slides with double-sided tape. Images were captured with a Leica S9i digital stereo microscope and analyzed using ImageJ. Wing length was measured from the alular notch to the intersection of the radius 3 vein and outer wing margin, disregarding fringe, as described previously [55].

The correlation between the measured wing lengths or time to pupation and alpha diversity of samples or relative abundance of ASVs of interest was calculated with a Generalized Estimating Equation Generalized Linear Model (GEE GLM) and mother/cohort as a cluster to account for differences in wing length due to genetic or other impacts of parentage. Success to adulthood, or the proportion of larvae in each isofemale line that reached adulthood within 18 days, was correlated with abundance of ASVs of interest using a binomial logistic regression. The number of larvae per cohort was included as a covariate in the models to account for its known influence on pupation timing (Additional file 4). Spearman’s rank-order correlation was used to investigate correlation in relative abundance between ASVs of interest.

For experiment 2, two development parameters were measured: proportion of eggs hatched and proportion of successful pupation events after 20 days. The correlation between offspring outcomes and relative abundance of genera of interest was calculated using a binomial logistic regression model weighted by number of eggs or number of larvae, respectively. *P*-values were adjusted for multiple comparisons using the Benjamini-Hochberg false discovery rate (FDR) procedure.

### Gnotobiotic larval assays to assess developmental effects of maternally-derived bacterial genera

In replicate C14 of Experiment 1, larvae did not progress past the first instar and larval death was observed. Six days after egg hatching, water from C14 was plated on Reasoner’s 2A Agar (R2A). Four bacterial isolates were isolated based on divergent colony morphology and stored in 20% glycerol stocks at −80 °C. DNA was extracted from an overnight culture and sent for Sanger sequencing with standard 16S rRNA gene primers (27F and 1492R) [56] at the DNA Sequencing Facility at the UW-Madison (Madison, WI USA). Larval development assays were conducted as previously described [7]. In short, axenic mosquito larvae were generated by successive rinses in 70% ethanol, 2% bleach, and 70% ethanol. Ten larvae were moved to each well of a six-well plate, and bacterial cultures were added to sterile larval water at a concentration of 5 μL of OD_600_ = 1 overnight culture per 5 mL of sterile water. For gnotobiotic coculture conditions, 1.25 μL of each OD_600_ = 1 overnight culture were added to 5 mL of sterile water, with the exception of strain HN178, for which 5 μL was added. To investigate the effect of individual isolates alongside a complex microbial community, 5 μL of OD_600_ = 1 overnight was inoculated along with 5 μL of cryopreserved conventionally reared (CR), insectary-derived larval rearing water. Larvae were fed a sterile rat chow diet as previously described and observed until pupation [7].

For a subset of maternally-derived genera associated with offspring developmental outcomes, experiments were also performed at the UW-Madison examine context-dependent effects on larval development. Isolates of *Asaia*, *Elizabethkingia*, *Paenibacillus*, and *Stenotrophomonas* were selected from the Mosquito-Associated Isolate Collection (MosAIC; [32]) and introduced into larval development assays as described above. Each isolate was either inoculated individually into larval water or co-inoculated with an equal volume of CR larval rearing water. Development was monitored for 18 days. Pupation success was compared among treatment groups using a Chi-squared test followed by Bonferroni-adjusted pairwise Fisher’s Exact tests. Wing length and time to pupation were analyzed using Kruskal-Wallis rank-sum tests with *post-hoc* Bonferroni-corrected Dunn’s tests. A list of all bacterial isolates used in the present study is provided in Additional File 5.

## Results

### Adult female mosquitoes transmit viable bacteria during oviposition that support development of larval offspring to the adult stage

In both experiments, laboratory-reared gravid (egg-bearing) female mosquitoes (and, in the case of Experiment 2, field-caught gravid females) were placed in sterile containers and allowed to oviposit. Oviposited eggs were then reared under aseptic conditions to the adult stage. In all lines except for one, larval offspring from an individual mother successfully developed to the L3/L4 stages, confirming the presence of viable bacteria that promote larval development. Control tubes in Experiment 1, which contained sterile water and diet but no larvae, showed no microbial growth on LB agar plates and minimal bacterial colonization with next generation sequencing methods, suggesting that all bacteria in the system were maternally transmitted.

### Maternally transmitted bacteria dynamically assemble into communities that recapitulate maternal bacterial diversity in offspring

The Experiment 1 sequencing effort comprised 186 experimental samples: F0 mothers (*n* = 23), F1 larval water (*n* = 114), and F1 larval offspring (*n* = 28), plus an additional 21 control samples. Overall, this resulted in a total of 7,855,925 sequences with a median of 33,325 sequences per sample and 423 unique amplicon sequence variants (ASVs) after quality control and filtering. After rarefying to a depth of 8,599 and removing samples with fewer reads, all but one true sample remained with a total of 348 unique ASVs. Alpha diversity as measured by Shannon’s *H* index was highest in F1 larval water samples and lowest in F0 mother and F1 larval offspring samples (Fig 2a). Higher larval density correlated with higher diversity in corresponding F1 larval offspring samples (Additional file 6).

**Figure 2:**
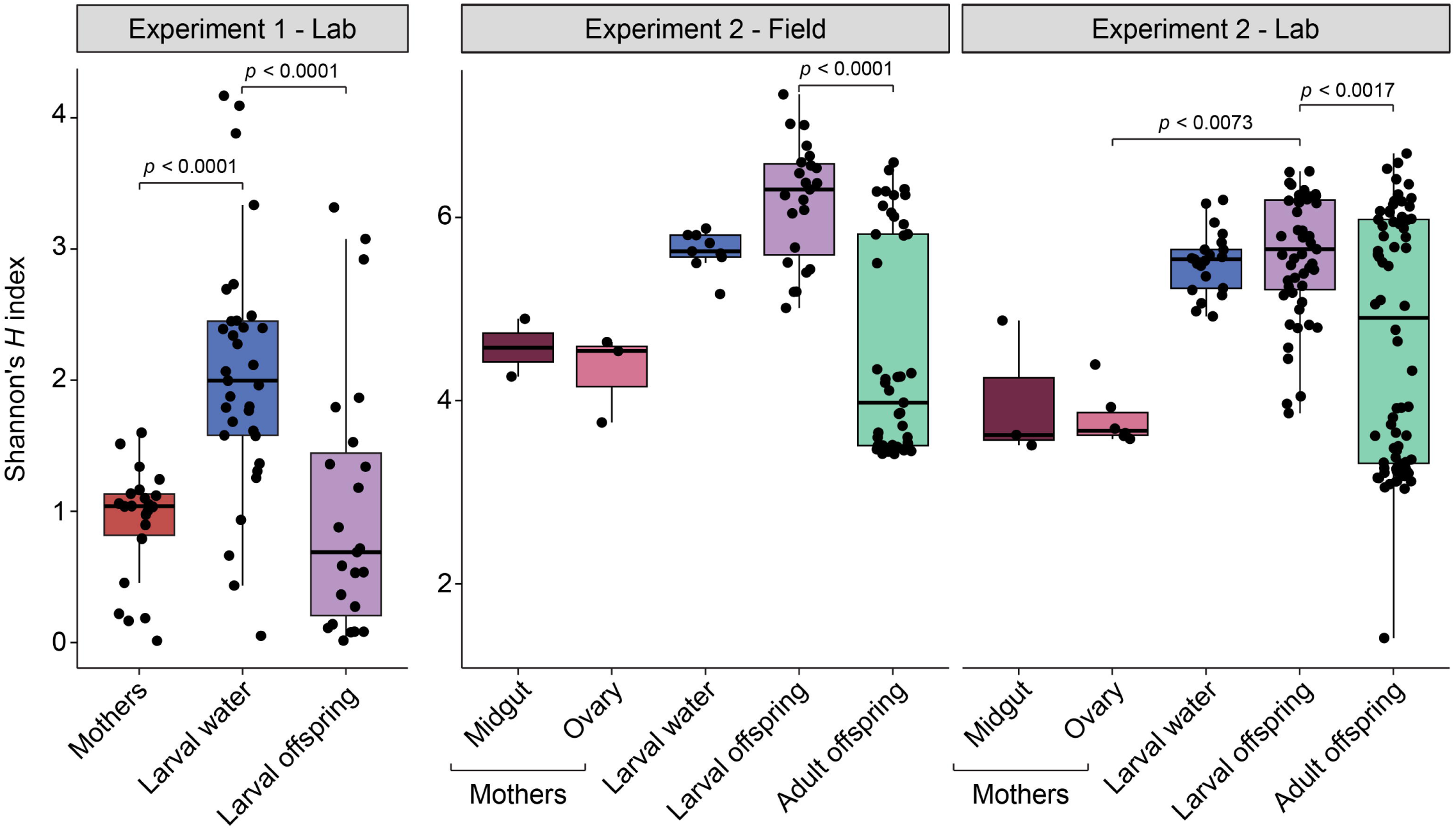
Differences in alpha diversity between sample types in Experiment 1 (*left panel*) and Experiment 2 (*right panel*), as measured by Shannon’s *H* Index. Box-and-whisker plots show median values with lower and upper edges of the box representing first and third quartiles. Significant differences were determined by Kruskal-Wallis Rank Sum followed by Dunn’s test for multiple comparisons. Experiment 2 data are faceted according to whether isofemale lines were derived from field-caught or laboratory-reared individuals.

The Experiment 2 sequencing effort comprised 268 experimental samples: F0 mother midguts (*n* = 10), F0 mother ovaries (*n* = 10), F1 larval water (*n* = 30), F1 larval offspring (*n* = 77), and F1 adult offspring (*n* = 141), plus an additional 7 control samples. Overall, this resulted in a total of 3,420,978 sequences with a median of 12,529 sequences per sample and 5,551 unique ASVs after quality control and filtering. After rarefying to a depth of 1,005 and removing samples with fewer reads, 237 true samples remained with a total of 3,412 unique ASVs. We first found that isofemale lines generated from laboratory-reared or field-caught mothers had no statistically significant difference in alpha diversity (Kruskal-Wallis rank-sum test; χ^2^ = 0.70981, *p* = 0.3995). Samples in Experiment 2 showed similar patterns of alpha diversity to those from Experiment 1, with F1 larval offspring samples harboring more diverse bacterial communities than F0 mother ovary samples (Fig. 2b). F1 adult offspring samples, which were sequenced in Experiment 2 but not Experiment 1, also harbored bacterial communities that were less diverse than F1 larval offspring samples but did not statistically differ from communities in F0 mother-derived samples (Fig. 2b).

Samples of larval water in Experiment 1 were collected at L1, L2-3, and L4 stages, as well as a final sample when the final larva pupated in each flask. Alpha diversity (Shannon’s *H* index) was not different across these four collection points (Kruskal-Wallis, χ^2^ = 4.9532, *p* = 0.1753). Ordination analysis of Bray-Curtis distances (a phylogenetically unaware metric) further revealed no differences in beta diversity between the same samples (Additional File 7, PERMANOVA, *F*_3,119_ = 1.1156, *p* = 0.332), although the unweighted Unifrac metric, which takes into account the phylogenetic relationships among the bacterial taxa in each sample, revealed that L1 larval water samples were distinct from other samples (Additional File 7, PERMANOVA, *F*_3,119_ > 2.48, *p* < 0.05). One ASV was more abundant in L1 larval water samples, which was an *Asaia* with reads comprising less than 1% of reads in most L1 water samples and nearly absent from all other time points (Additional File 8, aldex.kw, p< 0.05). L1 and L4 larval offspring formed distinct clusters by the Bray Curtis and unweighted Unifrac metrics (PERMANOVA *F*_1,28_ > 4.2, *p* < 0.01). Larval offspring collected at the L1 were more diverse (higher Shannon’s *H* Index) than L4 larvae (Additional File 10, Welch’s t-test, *t* = 4.5616). However, differential abundance analysis did not uncover any taxa significantly associated with L1 or L4 larvae. For all analyses that follow, Experiment 1 water or larval offspring samples were merged within a given isofemale line to maximize the number of isofemale lines with representative water and larval samples.

Ordination analysis of beta diversity using the unweighted Unifrac and Bray-Curtis distance metrics supported clustering of Experiment 1 samples by type, with F0 mothers, F1 larval water, and F1 larval offspring samples having distinct microbial signatures after accounting for significant batch effects (Additional File 11, PERMANOVA sample type *F*_2,72_ = 19.55*, p < 0.001*; batch *F*_2,72_ = 10.46*, p < 0.001*) (Fig. 3), though there were differences in dispersion between groups (Additional file 11). To generate gravid females for Experiment 1, three cohorts (batches) were reared in parallel from the same egg sheet. Differential abundance analysis with ALDEx2 revealed that larval water and larval offspring samples descended from mother in cohort A had higher *Elizabethkingia* abundance, often composing > 95% of reads in a given larval offspring sample (Fig. 7, Additional file 12). *Cedecea* was more common in descendants from cage B and C, while *Acinetobacter* was associated with cage C, suggesting that stochastic microbiota assembly within a generation can impact community composition of the subsequent generation.

**Figure 3:**
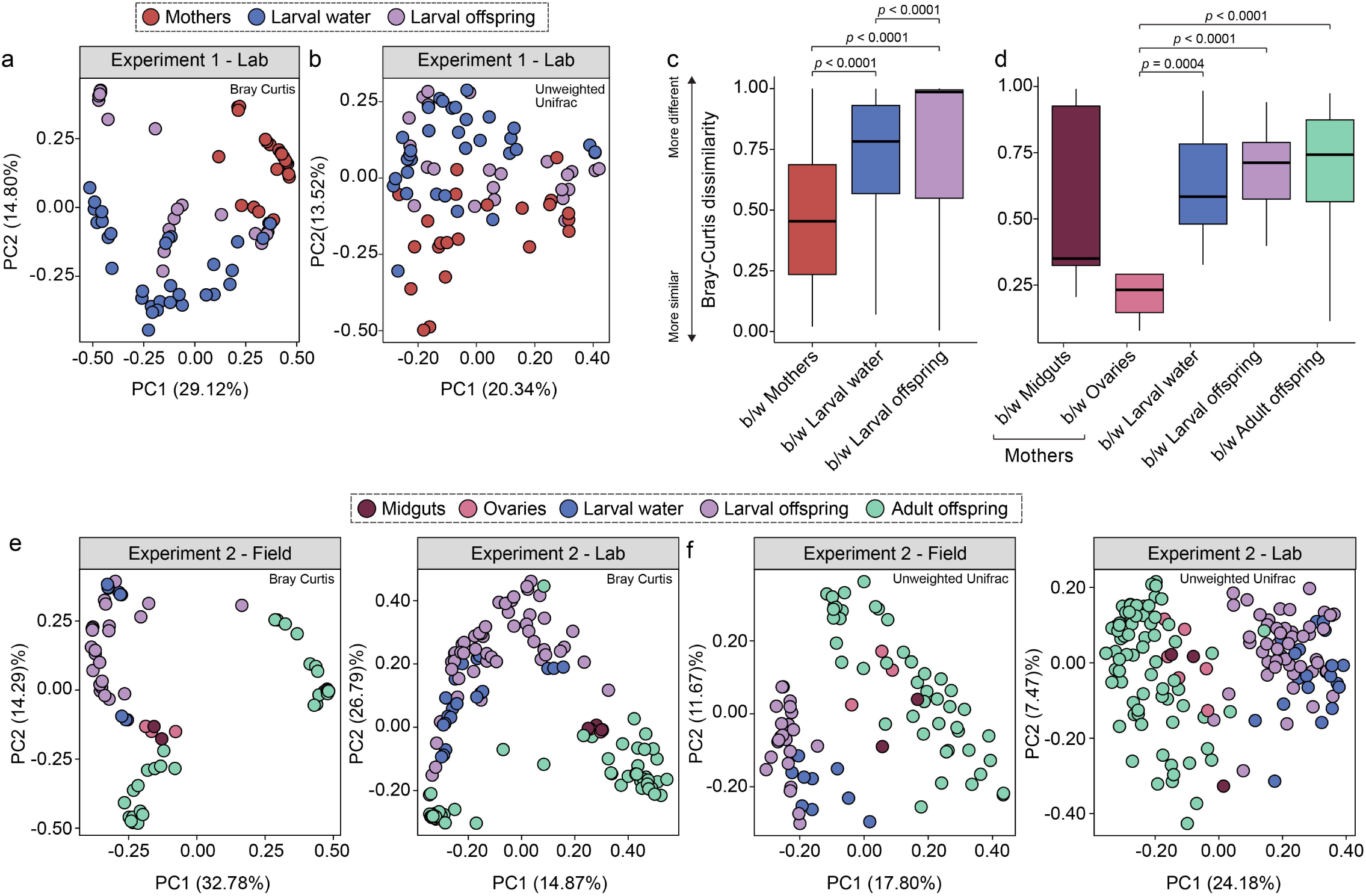
Principal coordinate analysis of differences in beta diversity among Experiment 1 (a, b) and Experiment 2 (e, f) samples, using the unweighted Bray-Curtis (a, f) and the unweighted Unifrac dissimilarity (b, e) indices. In Experiment 1, both indices support significant clustering by sample type (mothers, larval water, larval offspring) (Permutational Multivariate Analysis of Variance (PERMANOVA), p < 0.001, Additional File 10). When comparing differences between sample types for the laboratory-derived isofemale lines generated in Experiment 2, both the Bray-Curtis and Unweighted Unifrac dissimilarity index showed statistically significant clustering (PERMANOVA, p < 0.01, Additional File 12), except for midguts and ovaries, which clustered together. Using the unweighted Unifrac dissimilarity index in field-derived isofemale lines, there was also significant clustering by sample type (PERMANOVA, p < 0.01, Additional File 12), but ovaries and midguts clustered together with adult offspring. (c, d) Bray-Curtis dissimilarity between pairs of samples of the same sample type from Experiment 1 (c) and Experiment 2 (d), inclusive of both within-isofemale line pairs and between-isofemale line pairs. Box-and-whisker plots show median values with lower and upper edges of the box representing first and third quartiles. Significant differences were determined by Kruskal-Wallis Rank Sum followed by Dunn’s test for multiple comparisons.

Experiment 2 samples also generally clustered by type using both beta diversity metrics, with F0 mother-derived samples, F1 larval water, F1 larval offspring, and F1 adult offspring having distinct microbial signatures (Fig. 3, Additional File 13, PERMANOVA *F*_4,150_ 10.78, *p* < 0.001). In field-derived isofemale lines, F0 mother midgut and ovary samples clustered together with the F1 adult offspring samples when analyzed using the unweighted Unifrac dissimilarity index. This metric considers the phylogenetic relatedness of sample ASVs, suggesting that adult offspring communities recapitulate the diversity found in the previous generation. However, the same sample groups clustered separately when analyzed using the “phylogeny-unaware” Bray-Curtis dissimilarity index. (Fig. 3b; Additional File 13). There were statistically significant differences in dispersion, with more differences emerging among samples generated from laboratory-derived than field-collected isofemale lines, likely due to the larger sample size of the laboratory-derived group. F1 adult offspring generally showed the highest levels of dispersion (Additional File 13).

Differences in beta diversity within each sample type in Experiment 1, measured as average Bray-Curtis dissimilarity, were lowest amongst mother samples, and were notably higher within larval and water samples (Fig. 3c, Kruskal-Wallis Rank Sum test and Dunn’s test). Similarly, F1 larval water, larval offspring, and adult offspring all had significantly higher dissimilarity scores than F0 ovary samples in Experiment 2, suggesting that offspring communities vary more than the population of female mosquitoes that they originated from (Fig. 3d, Additional File 14). Despite the high dissimilarity within each offspring sample type, bacterial communities in samples derived from the same isofemale line had shared microbial signatures. Experiment 1 water and larval samples derived from the same isofemale line were more similar than water and larval samples from unrelated isofemale lines (Fig. 4b). In Experiment 2, Bray-Curtis dissimilarity was used to compare relationships within an isofemale line versus between unrelated lines, giving insight into the degree of intra-line variation compared to cross-line variation. In all sample types, regardless of whether the lines were lab- or field-derived, samples within an isofemale line were more similar to each other than to samples from other isofemale lines (Fig. 4d,e).

**Figure 4:**
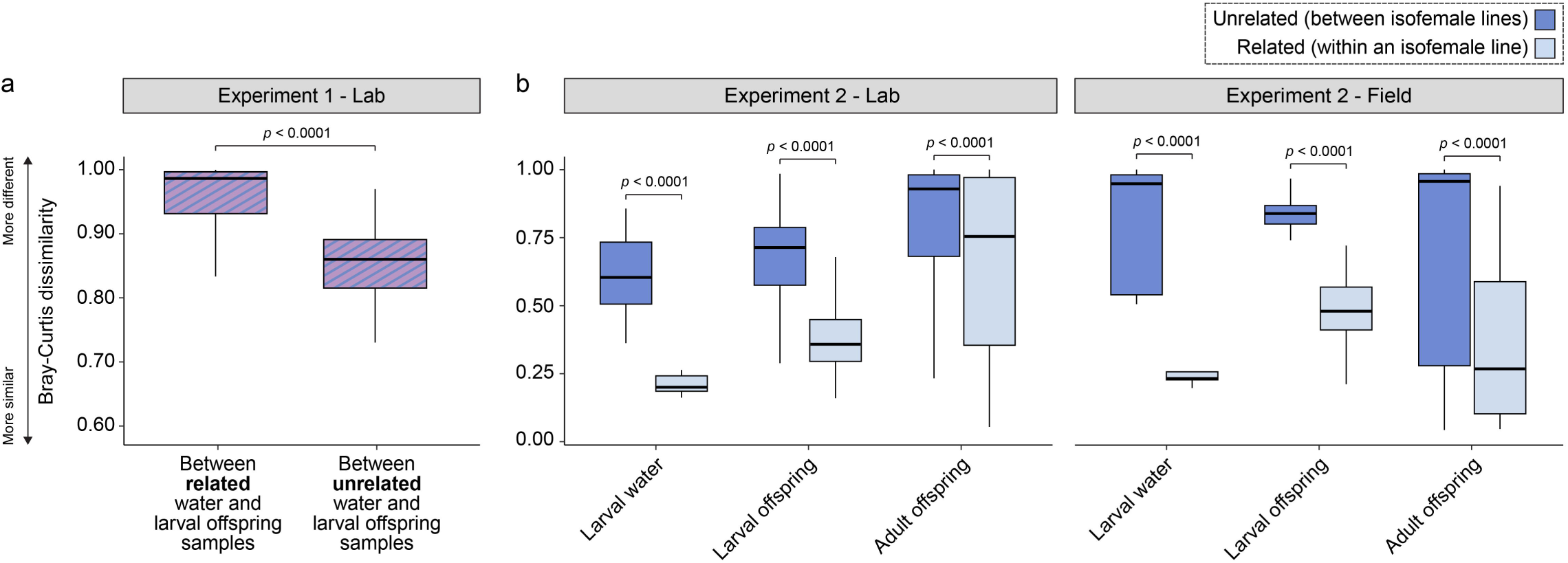
Bray-Curtis dissimilarity between pairs of samples of the same or different sample type from both experiments, estimated using the same distance matrices as in Figure 3. (a) Microbial community dissimilarity between pairs of water and larval offspring samples from the same isofemale line (related) or different isofemale lines (unrelated) in Experiment 1. (b) Dissimilarity between pairs of samples from related or unrelated isofemale lines in Experiment 2. Box-and-whisker plots show median values with lower and upper edges of the box representing first and third quartiles. Significant differences were determined by Welch’s t-test.

### Specific maternally transmitted taxa colonize and persistent in offspring

To gain a more detailed understanding of offspring-associated bacterial communities and their relationship to maternal communities, we identified bacterial taxa shared between mothers and offspring within the same isofemale line. Although offspring bacteria in these experiments were presumed to originate from mothers, examining ASVs shared with paired F0 samples revealed whether taxa dominating F0 samples also dominate F1 offspring. In Experiment 1, larval and water samples shared 1-10 ASVs with their paired mother samples, and these ASVs accounted for a variable proportion of offspring reads, ranging from 1% to 99.9% of reads in a given sample (Fig. 5a,b). While some ASVs were detected in offspring but not in the paired F0 female (Fig. 5), at the population level most reads could be traced back to mothers (99% in Experiment 1; 49% in Experiment 2). In Experiment 2, adult samples shared a higher percentage of reads with mothers than did larval samples (Additional File 15).

**Figure 5.**
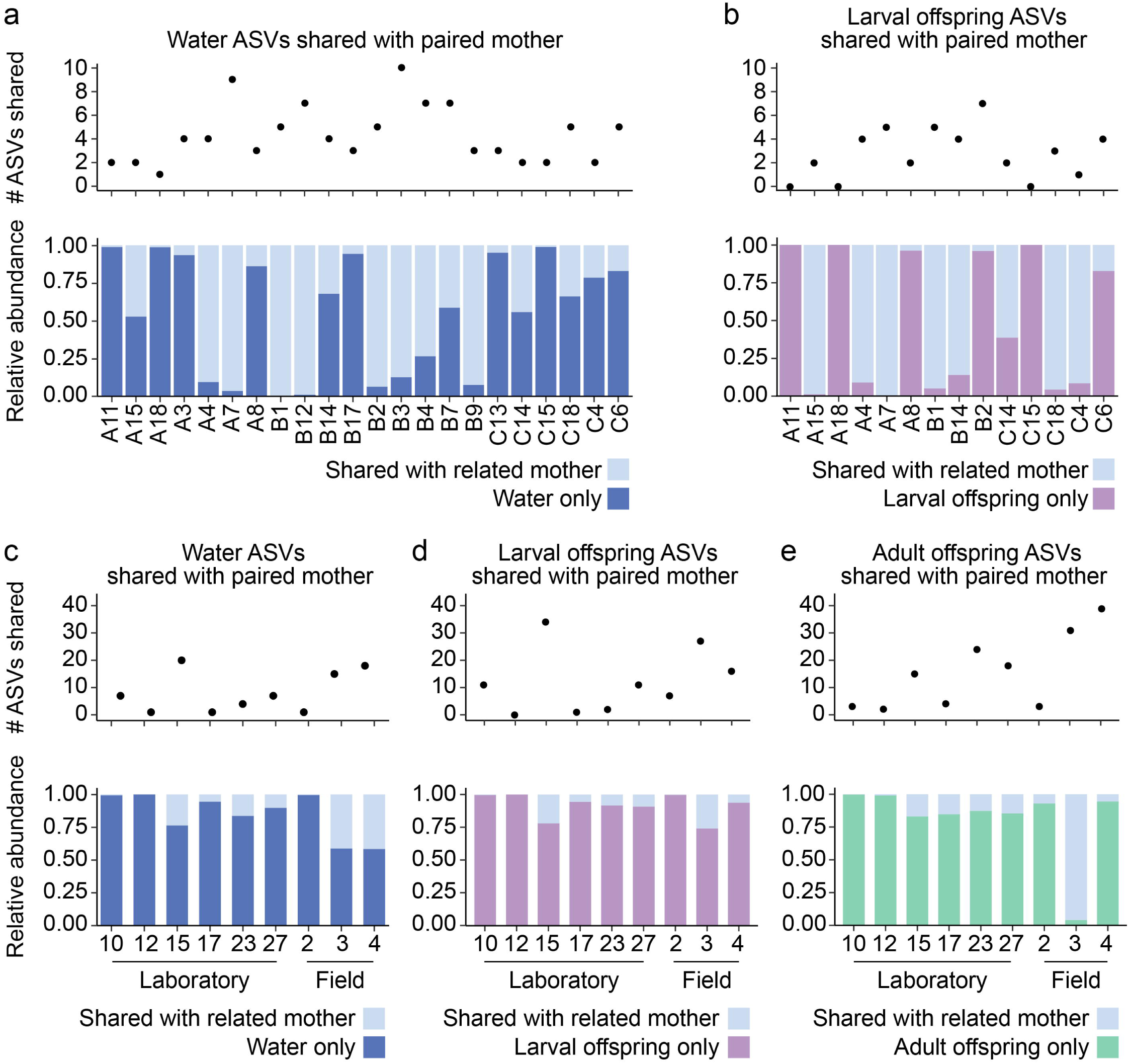
Number of amplicon sequencing variants (ASVs) shared between mothers (F0 samples) and related offspring (F1) samples, along with the relative abundance of shared versus unique ASVs in each sample type. Shared ASVs were defined as ASVs that were present (> 0 reads) in both the F0 mother sample and at least one F1 sample of a given sample type. Data for water (a, c), larval offspring (b, d), and adult offspring (e) for Experiment 1 (a,b) and Experiment 2 (c,d,e) are presented, pooled by sample type within an isofemale line to display the total proportion of reads shared with mothers regardless of variation between individuals.

The paired nature of these data allowed for tracking ASVs from a mother sample to subsequent water, larval, or adult samples. A successful transmission event was defined as detection of an ASV in a mother sample and at least one paired F1 sample of a given type. Considering Experiment 1, there were 15 ASVs that transmitted to water samples in more than one isofemale line (Fig 6), and 16 ASVs persisted to the larval stage across multiple isofemale lines. *Cedecea*, *Asaia*, *Elizabethkingia,* and an unclassified *Enterobacteriaceae* ASV were the most commonly transmitted ASVs (Figure 6a). The 25 most commonly transmitted ASVs in Experiment 2 were strikingly consistent between water, larval, and F1 adult samples, and almost entirely composed of *Burkholderia-Caballeronia-Paraburkholderia*, unclassified *Enterobacteriaceae*, and unclassified *Halomonadaceae*. *Paenibacillus* ASVs were commonly transmitted in laboratory-derived, but not field-derived, isofemale lines (Fig. 6b,c). We also considered which ASVs in F0 females were least likely to transmit to offspring. Of 198 ASVs in mother samples in Experiment 1, 133 were not transmitted to any F1 samples; however, these ASVs were rare (<5% of reads in a given F0 female), and only 22 were detected in more than one mother sample (Additional file 16). In Experiment 2, 133 out of 289 ASVs were not detected in any offspring. Only two of these ASVs were found in more than one mother sample, and all of these ASVs were rare (<5% of reads in a given F0 female) (Additional File 16).

**Figure 6.**
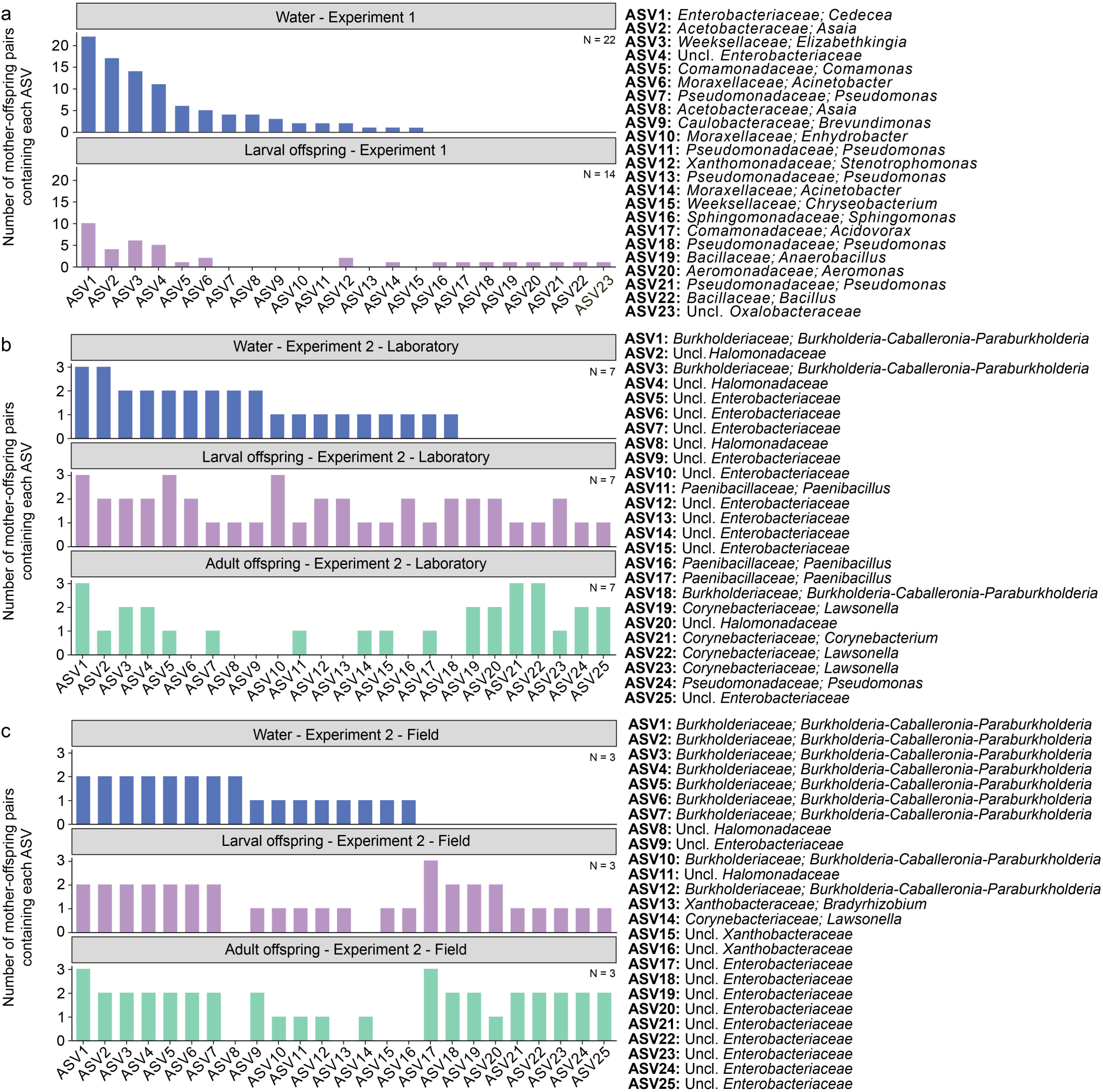
Frequency plots of the ASVs most often transmitted from mothers (F0) to related offspring (F1) samples in Experiment 1 (a), Experiment 2 laboratory-derived lines (b), and Experiment 2 field-derived lines. Transmission was defined as the detection of the same ASV in a given F0 mother and at least one related F1 offspring sample. ASV taxonomy, at the genus (or family) level, is provided on the right.

### Taxonomic composition of the mosquito microbiota depends on life stage and co-occurring taxa

Consistent with previous mosquito studies, the vast majority of reads from both Experiment 1 and Experiment 2 (83% and 69%, respectively) represented members of genera within the bacterial phylum Proteobacteria that varied in relative abundance between isofemale lines and life stages. In Experiment 1, we found low diversity in mother samples, which were dominated by *Asaia* and *Cedecea* (99% of reads) (Fig. 7). Despite the simple community detected in our single-mosquito extractions, offspring larvae and their water contained microbial communities that varied greatly from the paired mother sample and varied greatly from other larval samples. Larval samples were dominated by five genera that represented >75% of reads: *Elizabethkingia* (42%), *Cedecea* (20%), *Chryseobacterium* (9%), *Methylobacterium*-*Methylorubrum* (7%) *Stenotophomonas* (4%), and *Acinetobacter* (3%). Larval water samples were composed of mostly unclassified *Enterobacteriaceae* (25%), Cedecea (22%), *Elizabethkingia* (11%), *Comamonas* (8%), and *Brevundimonas* (7%). Differential abundance analysis with ALDEx2 identified two *Asaia* ASVs as positively associated with mother samples; while both of these ASVs were highly abundant in mother samples, they were low abundance in larval water and larval offspring samples (Additional File 17), which is consistent with work that demonstrated limited persistence of *Asaia* in larvae [57]. In contrast, a *Brevundimonas* and an unclassified *Enterobacteriaceae* ASV were positively associated with water but were rare in F0 female samples (Additional File 17). *Elizabethkingia* was highly abundant in both larval water and larval offspring samples (Additional File 17).

**Figure 7:**
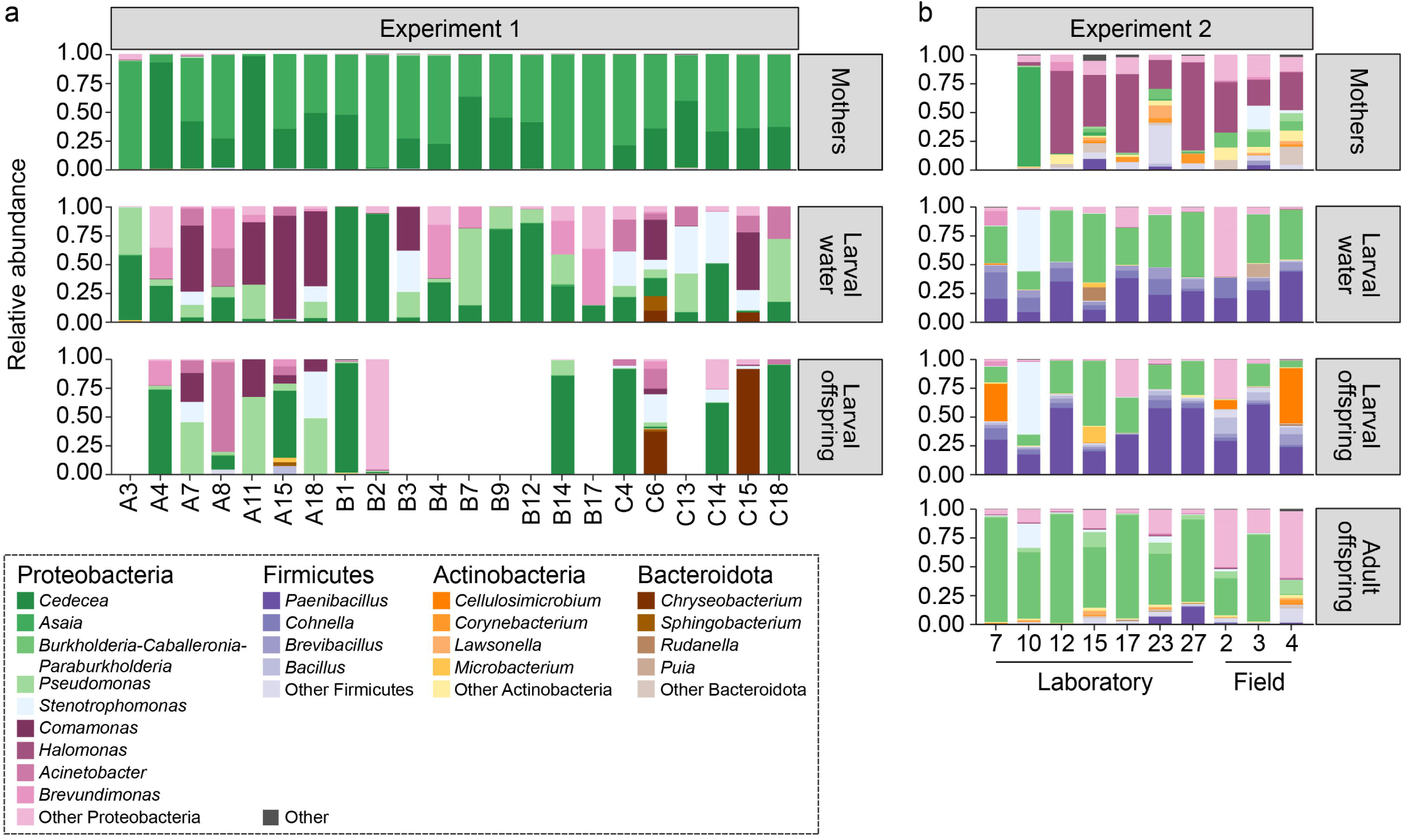
Stacked bar plots showing relative abundance plots of each bacterial genus within each isofemale line for Experiment 1 (a) and Experiment 2 (b). Each isofemale line resulted in numerous larval water and offspring samples, therefore the average relative abundances were calculated for each sample type. Experiment 2 mothers comprised midgut and ovaries samples; these are combined into one representative sample in the plots. Fully blank bars represent samples that did not pass sequencing quality control thresholds.

In addition to patterns of association with certain sample types, we found that certain ASVs exhibited patterns of co-occurrence or exclusion. In Experiment 1, *Elizabethkingia* exhibited exclusionary trends with the majority of abundant taxa in larvae, while the same ASV had neutral or positive correlations with most taxa in mother samples (Additional File 16). Similarly, a *Cedecea* ASV was negatively correlated with most other abundant ASVs in larval water (Additional File 18). Interestingly, there was a strong exclusionary trend between two *Asaia* ASVs in mother samples (ρ = −0.73396). Furthering the evidence for strain-specific interactions within mosquito microbiota, five different *Pseudomonas* ASVs were detected in larval water, each with a unique pattern of co-occurrence with other dominant ASVs in larvae. One of the *Pseudomonas* ASVs had a strong negative correlation with *Acinetobacter* in water samples (ρ = −0.6437). This same pair of ASVs had only weak negative correlation in larval offspring (ρ = −0.223) and was positively correlated in mother samples (ρ = 0.3505), suggesting host life stage affects not only the composition of microbial communities, but also the interactions between members of the community.

Experiment 2 also sought to identify bacterial communities that were passed from a mother to her offspring. In this experiment, the microbiota of the mothers’ midguts and ovaries were characterized and associated taxa were then traced through the larval water, larval offspring, and adult offspring samples. Similar to Experiment 1, here we saw considerable variation across isofemale lines and life stages, even when mother mosquitoes originated from the same laboratory colony and generation. Members of the Proteobacteria were present amongst all sample types (Figure 7b). Some genera were associated with particular sample types; *Halomonas* and unclassified *Halomonadaceae* were more common in midgut and ovary samples, *Paenibacillus* was more common in larval and larval water samples, and the genera *Pseudomonas, Aquabacterium*, and unclassified *Enterobacteriaceae* were most common in adult offspring (Additional File 19). The genus *Burkholderia-Caballeronia-Paraburkholderia* was abundant in all F1 sample types but rare in F0 mother samples. Additionally, spore-forming bacteria from the class Bacilli (*Paenibacillus*, *Brevibacillus*, *Bacillus*, *Lysinibacillus*, and *Domibacillus*) were more common in F1 samples (Additional file 19). Comparing adult offspring based on isofemale line origin, the majority of adult offspring of field-caught mothers harbored a considerable proportion of unclassified ASVs that belonged to the *Enterobacteriaceae* family, whereas adult offspring from lab-derived isofemale lines harbored more *Pseudomonas*, *Burkholderia-Caballeronia-Paraburkholderia,* and Serratia ASVs (Additional File 20). Within each taxonomic group, there were usually multiple ASVs present in each sample. These patterns of co-occurring ASVs within individual samples of particular sample types prompted us to further investigate positive or negative correlations between ASVs which could be suggestive of mutualistic or competitive bacterial interactions. Positive correlations within the unclassified *Enterobacteriaceae* and within the genus *Burkholderia-Caballeronia-Paraburkholderia* were seen in all sample types (Additional File 21-24). Larval offspring and water from laboratory-derived isofemale lines had common trends that did not appear in other sample types, including negative interactions between *Stenotrophomonas* and *Burkholderia-Caballeronia-Paraburkholderia* (−0.61 < ρ < −0.24) and a negative correlation between *Stenotrophomonas and Paenibacillus* (Additional file 21-22).

### Egg storage affects offspring microbial communities

All eggs from Experiment 2 were stored for three months under ambient insectary conditions prior to hatching, a practice which is common for *Ae. aegypti* colony maintenance [58, 59]. In Experiment 1, offspring originating from isofemale lines were floated in water within 24 hr of oviposition, and three population level “insectary control” lines were generated by using a desiccated egg sheet originating from the same population as the isofemale lines. Alongside methodological differences in larval diet and photoperiod between experiments, egg storage conditions may have affected resulting offspring bacterial communities. Notably, the L1 larval samples from the desiccated insectary control lines were dominated by *Paenibacillus* (Additional File 10), a genus of endospore-forming bacilli also common in larval and water samples in Experiment 2 (Additional File 19), though Experiment 1 L4 larval samples from insectary control lines converged toward the L4 larvae derived from isofemale lines (Additional File 10). Interestingly, water samples from the insectary control lines in Experiment 1 had higher alpha diversity than water from isofemale lines (Additional file 8, Welch’s t-test, t = 13, *p* < 0.001), suggesting that more diverse consortia of vertically transmitted microbes are deposited by a population of laboratory-reared insects compared to individual females from the same population.

### Dominant bacterial genera correlate with diverging host development outcomes

*Elizabethkingia* was a dominant member of the *Ae. aegypti* microbiome in Experiment 1, and its abundance correlated with positive offspring development outcomes. Isofemale lines with high *Elizabethkingia* abundance in water or larvae samples correlated with faster development (Additional file 25, GEE GLM, *p* = 0.0014, *p* < 0.0001), and larvae with higher relative abundances of *Elizabethkingia* also had higher proportions of success to adulthood (Additional File 25, binomial logistic regression, *p* = 0.011).

*Asaia*, previously reported to have a beneficial effect on larval development [57, 60], correlated with higher success to adulthood (Additional File 26, binomial logistic regression, *p* = 0.02) and larger wing size (GEE GLM, *p* = 0.008) when abundant in larvae. *Cedecea*, in contrast, correlated with slower offspring development when abundant in water and larvae samples (Additional File 26, GEE GLM, *p* = 0.00073, *p* = 0.022). Interestingly, the opposite seemed to be true for these ASVs in mother samples. When *Asaia* was abundant in mothers, offspring development was slower (Additional File 27, GEE GLM, *p* = 0.023), while high abundance of *Cedecea* in F0 females correlated with faster offspring development (Additional File 27, GEE GLM, *p* = 0.02). Notably, in neither case did abundance of *Asaia* or *Cedecea* in F0 females correlate with relative abundance of that ASVs in larvae samples (Additional File 28). In fact, abundance of these taxa in water was more predictive of larval offspring microbial communities (Additional File 28), suggesting that taxa at higher abundance in maternal samples are not guaranteed the same dominance in offspring samples.

In support of the negative effects of *Cedecea* on larval development, we observed that larvae hatching from eggs oviposited by a single female in Experiment 1 (C14) all died at the L1 stage without molting despite viable bacteria being readily recovered from larvae on LB agar plates. Further investigation confirmed that this killing phenotype was likely caused by an overgrowth of one strain of *Cedecea* (ASV d70bfa3628447052c56e1ff9d720c5c2, or *Cedecea_*d70) isolated from this treatment, a phenotype which did not occur when *Cedecea* was co-inoculated with a stock of larval rearing water containing the mixed community of bacteria present under conventional rearing conditions in our insectary (Additional file 29).

In Experiment 2, development impacts were investigated at the genus level. Higher relative abundance of *Asaia* in mothers correlated with a higher proportion of successful pupation (Additional file 30, binomial logistic regression, β = 23.1, *p* < 0.0001), while *Halomonas* was negatively correlated with offspring pupation (β = −27.58, *p* < 0.0001). *Burkholderia-Caballeronia-Paraburkholderia*, which was highly abundant in adult samples (Figure 7), had negative correlations with pupation success when present in larvae or water samples (β < −2.5, *p* < 0.001), while *Stenotrophomonas* in water and larvae positively correlated with pupation success (β > 2.9, *p* < 0.001). Interestingly, *Paenibacillus* abundance in water negatively correlated with pupation success (β = −2.44, *p* = 0.011) but not larval samples despite its prevalence in both sample types.

To further validate these correlations, we conducted gnotobiotic larval assays using a subset of taxa associated with developmental outcomes in Experiments 1 and 2. These assays, performed at the UW-Madison, allowed us to directly test the effects of individual isolates under controlled conditions. Axenic larvae inoculated with *Paenibacillus* rarely progressed beyond the first instar, and those that did were significantly delayed compared to axenic larvae reinoculated with conventional rearing water (Additional File 31). In contrast, taxa previously linked to positive outcomes—*Elizabethkingia* and *Asaia*—showed content-dependent effects: while both were associated with beneficial outcomes in maternally transmitted consortia, larvae inoculated with *Elizabethkingia* alone experienced slightly delayed pupation, and those inoculated with *Asaia* had reduced pupation success and smaller wings. Similarly, *Stenotrophomonas*, which correlated with pupation success in community-level analyses, caused delayed pupation, reduced wing size, and a slight but significant decrease in pupation success when tested in isolation. Notably, co-inoculation of these isolates with cryopreserved conventional rearing water restored normal development and adult wing length, underscoring the importance of microbial community context in shaping developmental trajectories.

## Discussion

In this work, we surveyed bacteria passed from laboratory- and field-derived female *Ae. aegypti* mosquitoes to offspring in a system free from external microbial inputs. We demonstrate that female mosquitoes deposit bacteria during oviposition capable of supporting offspring development. Offspring bacterial communities were found to vary greatly even when mothers are derived from the same insectary population and further show considerable variation in individual progeny of the same mother, potentially contributing to the natural variability of the microbiome seen in both lab and field mosquitoes. We identify key taxa detected across life stages and between data sets that correlate with different larval development outcomes, suggesting that variation in maternally derived bacteria may influence mosquito fitness and by extension human pathogen transmission [61, 62].

Vertical transmission of individual bacterial strains in mosquitoes has been demonstrated [11, 29, 30], but understanding of transmission of microbiota at the community level remains sparse. The life cycle of a holometabolous insect presents a challenge for vertical transmission. Mosquitoes have aquatic and terrestrial life stages, and the gut lining is shed during metamorphosis. As such, members of the gut microbiome are continually exchanged between the mosquito and larval habitat, are acquired through feeding on different sources across the mosquito life cycle, and are exposed to different selection pressures, resulting in a dynamic microbiome community. For this reason, the gut microbiota of mosquitoes is not classically considered a vertically transmitted system. However, we show that in a closed system, bacteria deposited by laboratory-reared and wild-caught female mosquitoes are sufficient to support development of offspring to adulthood. Potential mechanisms of maternal contribution include reproductive tract bacteria smeared on egg surfaces during oviposition, deposition of gut bacteria from defecation, and surface-associated microbes transferred via physical contact [27]. These maternally transmitted microbes are likely to interact with and compete with microbes present in larval water. In our experiments, female mosquitoes laid eggs in sterile water, but in their native habitat, mosquitoes oviposit in water already colonized by microbes and shaped by environmental factors. Field larval habitats differ not only in microbial composition but also in physiochemical conditions, such as pH, organic matter, and nutrient availability, that can influence microbial persistence, transfer, and favor certain taxa over others. Oviposition itself can also alter the microbial and chemical profile of water [31], introducing maternal microbes and changing conditions in ways that may affect community assembly. Additional work is required to determine how maternal microbes compete with or are facilitated by these environmental conditions to colonize, persist, and influence host development.

Interestingly, bacterial diversity was higher in larval water than in corresponding lab-derived mother samples. While we acknowledge the limitations of sequencing low biomass samples of a single mosquito midgut, ovary, or abdomen, our closed rearing system ensured that any bacteria found in larvae was present in mothers, albeit at low levels, even if not detected in maternal sequencing data. Thus, we show that high abundance in mothers is not a prerequisite for passage of bacteria to the next generation. In fact, adult offspring communities resembled midgut and ovary samples more than water or larvae samples. This suggests that the differences between larval and adult diet, physiology, and environment have a strong effect on bacterial community composition even when the pool of available bacteria is limited to maternal inputs. Furthermore, we observed variation in microbiota composition among individual offspring from a single mother, which differed between life stages and between lab- and field-derived isofemale lines. Despite mothers from Experiment 1 being reared in three parallel cohorts originating from the same egg sheet, we saw large differences in offspring microbial community between maternal cohorts. Offspring from cohort A were dominated by *Elizabethkingia,* cohorts B and C were dominated by *Cedecea*, and *Acinetobacter* was abundant in cohort C but nearly absent in other cohorts. This demonstrates the stochastic nature of microbiome assembly and underscores the need for further research to understand the factors influencing microbiome assembly in mosquitoes.

While vertical transmission of microbiota may occur in all mosquito species, here we chose to use *Ae. aegypti* due to ease of experimental manipulation and its status as a vector of human pathogenic viruses. Vertical transmission of single bacterial isolates has been demonstrated in other species of mosquitoes and suggests that complex bacterial communities are transmitted in other species as well, though further work is required to determine how vertical transmission differs in other key vector species [11, 29, 30, 63]. Unlike vector species from the genera *Anopheles* and *Culex*, the eggs of *Ae. aegypti* are desiccation resistant. Both the environmental conditions and composition of the egg surface impact which bacteria survive long enough to colonize offspring in natural environments, so the particular challenges to bacterial transmission are varied between species. Our experiment provides insights into the effect of egg desiccation on vertical transmission of microbial communities. In Experiment 2, all eggs were stored for three months before use. In this experiment, a smaller percentage of offspring reads belonged to ASVs also detected in mother samples, and larval samples were associated with endospore forming members of the class Bacilli. L1 larvae from desiccated “insectary control” eggs in Experiment 1 were also enriched in *Paenibacillus*, suggesting that consistent maternally transmitted bacteria survive egg desiccation and can in turn dominate larval and water communities. Indeed, *Paenibacillus* has been detected as a component of wild-collected mosquito microbiome, perhaps due to its ability to persist through periods of desiccation [62–64]. In the present study, *Paenibacillus* was correlated with negative development outcomes, which was supported by gnotobiotic larval development assays where 93% of larvae failed to progress past the first instar.

In accordance with previous studies, both experiments abundantly featured members of the *Enterobacteriaceae* in the mosquito microbiome [65, 66]. Common mosquito-associated genera such as *Serratia* and *Pseudomonas* were present in multiple samples across both experiments [67, 68]. We also found the community to be dynamic and variable across experiments and both across and within isofemale lines, with particular taxa only being seen in one experiment or condition. *Asaia*, previously reported to improve mosquito development outcomes [57, 60] played a key role in this study. In Experiment 1, its presence in larvae correlated with positive development outcomes, and in Experiment 2, mothers with more *Asaia* had offspring with faster time to pupation. Subsequent gnotobiotic larval development assays demonstrated that *Asaia* alone does not support pupation success, but inoculation alongside a diverse microbial consortium precludes any deleterious effects. Interestingly, the relative abundance of *Asaia* was quite low in Experiment 1 larval offspring, but multiple studies have noted a positive effect of adding *Asaia* to developing larvae even when levels quickly become undetectable, suggesting that *Asaia* may affect larval development outcomes through modifications to bacterial community assembly [31, 57]. *Cedecea* was prominent in Experiment 1, which is consistent with its appearance in previous literature [28, 67, 69–71]. We found that *Cedecea* abundance in mothers is correlated with positive offspring development outcomes, but high *Cedecea* abundance in larvae and water correlated with negative outcomes. This negative correlation was confirmed by rearing gnotobiotic mosquitoes with *Cedecea* isolated from an isofemale line with high larval mortality; these monocolonized larvae died or arrested at the L1 stage. However, it is well known that bacteria do not act in isolation. In fact, *Cedecea* has been shown to inhibit *Serratia* colonization in mosquitoes [67]. In Experiment 1, higher relative abundance of *Asaia* in mothers correlated with longer time to pupation, perhaps due to exclusionary interactions with a *Cedecea* ASV. In this study, we found additional positive and negative correlations in bacterial abundance in this system that varied between life stages. Notably, in Experiment 1, two *Asaia* ASVs were negatively correlated only in mother samples. Coinfection with multiple *Asaia* strains occurs in *Anopheles* and *Aedes* mosquitoes [72], which has implications for selecting strains for paratransgenesis that can coexist with or displace native bacteria to achieve both reliable pathogen blocking and transmission across generations. In Experiment 2, the only correlations that persisted across all sample types were positive associations within members of the *Enterobacteriaceae* and within the genus *Burkholderia-Caballeronia-Paraburkholderia*. There were many more pairs of ASVs that showed correlations only in particular sample types. We saw a negative correlation between *Burkholderia-Caballeronia-Paraburkholderia* (previously *Burkholderia*) and *Pseudomonas* ASVs in adult offspring of laboratory-derived mothers only; both of these genera were previously noted as exclusionary players in network analyses [69]. The variations in correlation patterns in different sample types across both experiments, and in previous work, are likely influenced by diverse factors including external environmental conditions, conditions within the host and the presence/absence of additional community members. The contributions of these factors warrant further research if we are to fully understand the drivers of microbiome assembly and transmission.

In a publishing environment that favors novelty over repetition [73], it is rare to see results replicated in different labs. Here, experiments were independently conceived and conducted by two separate laboratories, and the analysis was conducted in tandem. This work highlights the benefit of collaborative science, as each group addressed the same core question but emphasized different aspects of mosquito biology. For example, both labs allowed female mosquitoes to oviposit under sterile conditions and traced the microbial community of her offspring. These experiments revealed similar trends in community composition and validation that mosquitoes can vertically transmit complex communities. Distinguishing the two approaches, Experiment 1 investigated the fitness effect of variable microbiota transmitted from mothers directly after oviposition, while Experiment 2 considered the differences in field- versus lab-derived mothers and included an additional sequencing group of F1 adults. Together, these combined studies provide a more holistic picture of vertical transmission. We acknowledge that methodological differences between the two experiments may influence microbiota composition and thus affect comparisons. Key differences include photoperiod (16:8 vs. 12:12 light:dark), larval diet (rat chow vs. fish flakes), and adult diet (5% vs. 10% sucrose). Diet at both life stages can strongly impact host immunity and microbiota composition [74–76]. Photoperiod influences mosquito behavior [77–79], though its effects on gut microbial communities remain unknown. Additionally, the sequenced 16S rRNA region differed (V4 vs. V3-V4), and rarefaction depths for alpha and beta diversity analyses varied due to differences in sequencing depth and ASV-level diversity. The longer region sequenced in Experiment 2 yielded more ASVs, reflecting higher taxonomic resolution. For these reasons, we did not perform quantitative comparisons between datasets and instead focused on genus-level similarities, using the same classification database for both studies.

Through the use of 16S rRNA gene amplicon sequencing data, this study assessed community composition and traced ASVs passed from female mosquitoes to offspring. Future work may investigate variation of transmission at the species or strain level, as different strains of bacteria have distinct functional capabilities. Particular bacterial genes or traits that enhance colonization and intergenerational persistence could be a crucial tool for enhancing efficacy of paratransgenesis efforts. Furthermore, the degree of stochasticity observed in microbiome assembly within isofemale lines reared in closed systems highlights the complexity of interacting factors governing microbiome assembly. Particularly, future work may address bacterial interactions within the microbiome. For bacteria employed to influence the pathogen transmission ability of mosquitoes, competition with other bacteria on the axes of nutrient availability, space, and transmission to future generations may have large consequences for human health. Further studies should consider the impact of variation in fidelity of vertical transmission of microbes in additional vector species or under differing conditions. Finally, a growing body of work highlights the role of fungi in the mosquito gut community [80–82]. Expanding vertical transmission to not only fungi, but also considering other microbiome members such as protists and insect-specific viruses [83–86] and determining the relative contributions of maternal versus environmental bacteria to the mosquito microbiome will reveal avenues to improve our use of microbes as vector control and shed light on the biology related to transmission of these microbes in mosquitoes.

## Conclusions

Taken together, our results suggest that female mosquitoes deposit complex bacterial communities that support offspring development. Adult mosquito offspring raised in a closed system harbor microbiota resembling that of their mother despite changes during the larval stage, and the variable composition of the inherited microbiome correlates with divergent mosquito development outcomes. This result lays the foundation for further research to examine vertical transmission in the mosquito microbiome, which will aid in manipulating mosquito microbiota for controlling mosquito-borne diseases.

## List of Abbreviations

ASV: amplicon sequence variant
PERMANOVA: permutational multivariate analysis of variance
PERMDISP: permutational multivariate analysis of dispersion
LVP: Liverpool strain of *Aedes aegypti*
GALV: Galveston strain of *Aedes aegypti*

## Declarations

### Ethics approval and consent to participate

Not applicable.

### Consent for publication

Not applicable.

### Availability of data and materials

Raw Illumina reads are available in the NCBI Sequence Read Archive ([80–82]) under BioProject IDs PRJNA1054601 and PRJNA1055422. Scripts used for analysis and figure generation are available in the following GitHub repository: https://github.com/kcoonlab/mosquito-vertical-transmission. All other data generated by this study are available as Supporting Information herein.

### Author contributions

HLN, KLC, MAS, and GLH conceived of and designed the experiments; HLN, KV, MAS, and SH performed the experiments; HLN and KLC carried out the data analysis, with early input from LEB; HLN and KLC wrote the manuscript, and MAS, LEB, and GLH provided comments during revision.

## Acknowledgements

We thank Dr. Lyric Bartholomay and the insectary staff at the UW-Madison and the UTMB for maintaining the *Ae. aegypti* Liverpool (LVP) and Galveston (GALV) laboratory colonies used for Experiment 1 and Experiment 2 in this study, respectively.

## Additional File Captions

**Additional File 1:** Supplemental methods figure detailing sampling scheme in Experiment 1 and Experiment 2. See Fig. 1 and relevant methods section for additional information.

**Additional File 2:** Rarefaction curve generated for Experiment 1 showing a plateauing of species richness at the rarefaction depth chosen for subsequent analyses (8599 reads).

**Additional File 3:** Rarefaction curves generated for Experiment 2 showing a plateauing of species richness at the rarefaction depth chosen for subsequent analyses (1005 reads).

**Additional File 4:** Larval density and fitness metric correlations Experiment 1. Variation in the number of larvae per experimental flask significantly correlated with development outcomes and is included as a factor in all subsequent models. Linear regression.

**Additional File 5:** List of bacterial strains used in the present study.

**Additional File 6:** Alpha diversity in Experiment 1 (Shannon’s H Index) in mother and water samples is predictive of alpha diversity of paired larval samples. Linear regression.

**Additional File 7:** Experiment 1 Permutational Multivariate Analysis of Variance (PERMANOVA) and Permutational Multivariate Analysis of Dispersion (PERMDISP) results (both Bray-Curtis and unweighted Unifrac) comparing beta diversity of isofemale line larval water between four sampling time points including early water (24 h after hatching) mid water (3 days after hatching), late water (collected on day of first pupation in each flask) and final (pupal) water (18 days after hatching).

**Additional File 8:** Comparison of microbial community in larval water at four sampling time points. (a) Relative abundance of bacterial genera in Experiment 1 larval water samples over four sampling time points. Amplicon Sequencing Variants (ASVs) are colored by genus. Isofemale lines (derived from a single ovipositing female) are represented by A, B, or C to denote the replicate cohort that mothers were derived from, and insectary control lines (derived from a standard egg sheet from the same population of insectary-reared *Aedes aegypti*, eggs stored for approximately 3 months before hatching) are represented by letters IC. (b) Alpha diversity of larval water samples does not differ by sampling time point as measured by Shannon’s H Index (Kruskal-Wallis, H = 4.9532, *p* = 0.1753). (c) Relative abundance of the only ASV significantly different between water sampling time points, classified as *Asaia*. This ASV is most abundant in L1 larval water (ALDEx2 Kruskal-Wallis, p < 0.001). (d) Alpha diversity of larval water samples is higher in insectary control lines (Welch’s t-test, p < 0.0001).

**Additional File 9:** Experiment 1 PERMANOVA and PERMDISP results (both Bray-Curtis and unweighted Unifrac) comparing beta diversity of isofemale line larval offspring between two sampling time points including L1 larval offspring (collected 24h after hatching) and L4 larval offspring (collected on the day of first pupation in each flask).

**Additional File 10:** Comparison of microbial communities in Experiment 1 larvae at two sampling time points, considering larvae from isofemale lines and from insectary population egg sheets stored under standard conditions (insectary control). (a) Relative abundance plot of Experiment 1 L1 versus L4 larval offspring samples from isofemale (A, B, C denoting replicate cohorts of mothers) and insectary control (IC) lines. Isofemale lines are derived from a single ovipositing female, and insectary control lines are derived from a standard egg sheet from a population of insectary-reared *Aedes aegypti*, eggs stored for approximately 3 months before hatching. ALDEx2 did not identify any differentially abundant ASVs between L1 and L4 larval samples. (b) Alpha diversity, represented by Shannon’s H Index, is higher in L1 larval offspring than L4 larval offspring (Welch’s t-test, t = 4.56, *p* = 0.0003)

**Additional File 11:** Experiment 1 PERMANOVA and PERMDISP results (both Bray-Curtis and unweighted Unifrac) comparing beta diversity between sample types.

**Additional File 12:** Relative abundance of genera in Experiment 1 identified as significantly differentially abundant in offspring samples (larval offspring and larval water) across maternal cohort as evaluated by ALDEx2 (aldex.kw, p < 0.05).

**Additional File 13:** Experiment 2 PERMANOVA and PERMDISP results (both Bray-Curtis and unweighted Unifrac) comparing beta diversity between sample types.

**Additional File 14.** Bray-Curtis dissimilarity boxplots for Experiment 2 demonstrate large differences between individuals within a given isofemale line. For larval water, larval offspring, and adult offspring samples, only within-isofemale line comparisons are considered. This highlights the differences between individuals in the F0 population (ovary samples) compared to differences between individuals in the F1 population (adult offspring). Significant differences were determined by Kruskal-Wallis and Dunn’s test with Bonferroni correction for multiple comparisons.

**Additional File 15.** Average relative abundance of F1 sample ASVs shared with at least one F0 mother sample in Experiment 1 (a) and Experiment 2 (b). Bars represent the standard error of the mean. No significant differences for Experiment 1 samples were detected (Mann-Whitney U test; *W* = 350, *p* = 0.4612), while Experiment 2 adult offspring samples harbored significantly higher relative abundances of shared ASVs than larval offspring samples (Kruskal-Wallis Rank Sum test followed by Dunn’s test for multiple comparisons; H = 18.569, *p* < 0.001).

**Additional File 16.** ASVs commonly lost. All ASVs found in mother samples were considered, and an ASV was considered “lost” if it was present in at least one mother sample but found in no offspring samples in the population. Lost ASVs that were found in more than two mother samples in Experiment 1 are displayed (a), and the fifteen lost ASVs with the highest read count in Experiment 2 are displayed (c). Relative abundance of mother ASVs in Experiment 1 (b) and Experiment 2 (d) are color-coded by whether each ASV was detected in any offspring samples (blue) or not detected in any offspring samples (red).

**Additional File 17.** Relative abundance of ASVs in Experiment 1 identified as significantly differentially abundant across sample types as evaluated by ALDEx2 (aldex.kw, p < 0.05).

**Additional File 18:** Correlation plots: Experiment 1. Spearman’s rank-order correlation of relative abundance of the 25 most abundant ASVs found in (a) mother, (b) larval offspring, or (c) larval water samples. Color represents Spearman’s rho (ρ), where positive (blue) numbers represent a positive co-occurrence (synergistic) and negative (red) numbers represent exclusionary (antagonistic) patterns of abundance. Three characters from the ASV code have been added to distinguish ASVs from within a genus.

**Additional File 19.** Relative abundance of genera in Experiment 2 identified as significantly differentially abundant across sample types as evaluated by ALDEx2 (aldex.kw, p < 0.05).

**Additional File 20.** Effect plot of ASVs in F1 adult samples in Experiment 2 differentially abundant between lab- and field-derived isofemale lines. ASVs identified as significantly differentially abundant by both t.test and effect parameters in ALDEx2 (p < 0.05) are highlighted in green, red, blue, or yellow. Axes represent the difference and dispersion of the relative abundance of each ASV between the two groups (Field- or Lab-derived F1 adult samples). Points are colored by taxonomy while dashed lines represent equal difference (between-group variation) and dispersion (within-group variation).

**Additional File 21:** Correlation plots: Experiment 2 showing correlation of relative abundance of common ASVs in ovary samples from laboratory- and field-derived mothers. Color represents Spearman’s rho, where positive numbers represent a positive co-occurrence (synergistic, blue) and negative numbers represent an exclusionary (antagonistic, red) pattern of abundance. Taxa descriptions are shown family and genus level where applicable.

**Additional File 22:** Correlation plots: Experiment 2 showing correlation of relative abundance of common ASVs in water samples from laboratory- and field-derived isofemale lines. Color represents Spearman’s rho, where positive numbers represent a positive co-occurrence (synergistic, blue) and negative numbers represent an exclusionary (antagonistic, red) pattern of abundance. Taxa descriptions are shown family and genus level where applicable.

**Additional File 23:** Correlation plots: Experiment 2 showing correlation of relative abundance of common ASVs in larval offspring from laboratory- and field-derived isofemale lines. Color represents Spearman’s rho, where positive numbers represent a positive co-occurrence (synergistic, blue) and negative numbers represent an exclusionary (antagonistic, red) pattern of abundance. Taxa descriptions are shown family and genus level where applicable.

**Additional File 24:** Correlation plots: Experiment 2 showing correlation of relative abundance of common ASVs in F1 adult offspring from laboratory- and field-derived isofemale lines. Color represents Spearman’s rho, where positive numbers represent a positive co-occurrence (synergistic, blue) and negative numbers represent an exclusionary (antagonistic, red) pattern of abundance. Taxa descriptions are shown family and genus level where applicable.

**Additional File 25:** Relative abundance of *Elizabethkingia*_b61 correlates with larval development outcomes in Experiment 1. (a) Higher relative abundance of *Elizabethkingia_*b61 in larval offspring correlates with (a) faster development as measured by days to pupation (Generalized estimating equation (GEE) GLM) and (b) higher proportion of survival to adulthood (binomial logistic regression). Higher relative abundance of *Elizabethkingia_*b61 in larval water also correlates with (c) faster development (GEE GLM)

**Additional File 26:** Higher relative abundance of *Asaia*_5aa in larvae is correlated with (a) higher proportion of larvae surviving to adulthood and (b) larger wing length in a given isofemale line in Experiment 1. However, when this *Asaia*_5aa abundance is higher in F0 mother samples, paired offspring have delayed development (c).

**Additional File 27:** Higher relative abundance of *Cedecea_*d70 in water or larvae is correlated with slower development time as measured by average days to pupation in a given isofemale line (GEE GLM) in Experiment 1, but higher abundance of the same ASV in mother samples is correlated with faster development time in given isofemale line (GEE GLM).

**Additional file 28:** Relative abundance of *Asaia*_5aa in larval water, but not mother samples, correlates with *Asaia*_5aa abundance in larvae in Experiment 1. Similarly, relative abundance of *Cedecea_*d70 in larval water, but not mother samples, correlates with *Cedecea_*d70 abundance in larvae (quasibinomial logistic regression).

**Additional File 29.** Proportion of larvae that successfully pupated by Day 9 of development, sample size on graph. (a) Larvae exposed to individual bacterial isolates isolated from C14 replicate of Experiment 1. HN176 and HN178 have a significantly lower pupation rate compared to CR (conventionally reared) larvae (Chi-Square, Pairwise Fisher’s Exact test p < 0.05). To test if the larval killing phenotype still occurred in a bacterial community, larvae were exposed to multiple conditions: C14_coisolates is an inoculum composed of all isolates tested in (a), “top4” is composed of 4 bacterial isolates from the Mosquito Associated Isolate Collection (MosAIC) with 100% 16S rRNA sequence identity to abundant ASVs detected in the Experiment 1 dataset, including MMO-72 (*Chryseobacterium*), MMO-130 (*Acinetobacter*), USMM111 (*Brevundimonas*), and MMO-42 (*Comamonas*). HN178 was added to these defined communities or to a mixed stock of microbes from conventionally reared (CR) mosquitoes. HN178 and top4_HN178 had significantly lower pupation rates than CR positive control (Chi-Square, Pairwise Fisher’s Exact test with Bonferroni correction, *p* < 0.05).

**Additional File 30**: Commonly detected genera in Experiment 2 correlate with F1 development outcomes. (a) Genera in mother samples that correlate with egg hatch rate. (b) Genera in mother samples that correlate with F1 pupation rate. (c) Genera in water samples that correlate with pupation rate. (d) Genera in larval offspring samples that correlate with pupation rate. Binomial logistic regression model weighted by number of eggs (a) or larvae (b-d) in each isofemale line.

**Additional File 31** Larval development in gnotobiotic assays. Bacterial isolates were selected from genera associated with offspring development outcomes in isofemale lines from Experiment 1 or Experiment 2. Isolates were obtained from the Mosquito Associated Isolate Collection (MosAIC) [31] or from a laboratory colony of *Aedes aegypti* (Liverpool strain) at the University of Wisconsin-Madison. (a) Days to pupation; treatments significantly different from conventionally reared (CR) group are indicated by an asterisk (Kruskal-Wallis rank-sum test followed by Dunn’s multiple comparisons, *p* < 0.05). (b) Wing length of newly emerged adults; analyzed using one-way ANOVA followed by Dunnett’s *post-hoc* test comparing each treatment to CR (*p* < 0.05). (c) Proportion of successful pupation by Day 18; sample sizes shown on the plot (Chi-square test with pairwise Fisher’s Exact tests and Bonferroni’s correction, *p* < 0.05).

**Additional File 32:** Sequence metadata for Experiment 1 containing sample id, metadata info, and total reads.

**Additional File 33:** Development data (days to pupation and wing length) for Experiment 1.

**Additional File 34:** Sequence metadata for Experiment 2 containing sample id, metadata info, and total reads.

**Additional File 35:** Development data for Experiment 2 by isofemale line.

