## Supplementary figures and images for "Vertical transmission of mosquito microbiota and its effects on offspring development"

### af_1.pdf

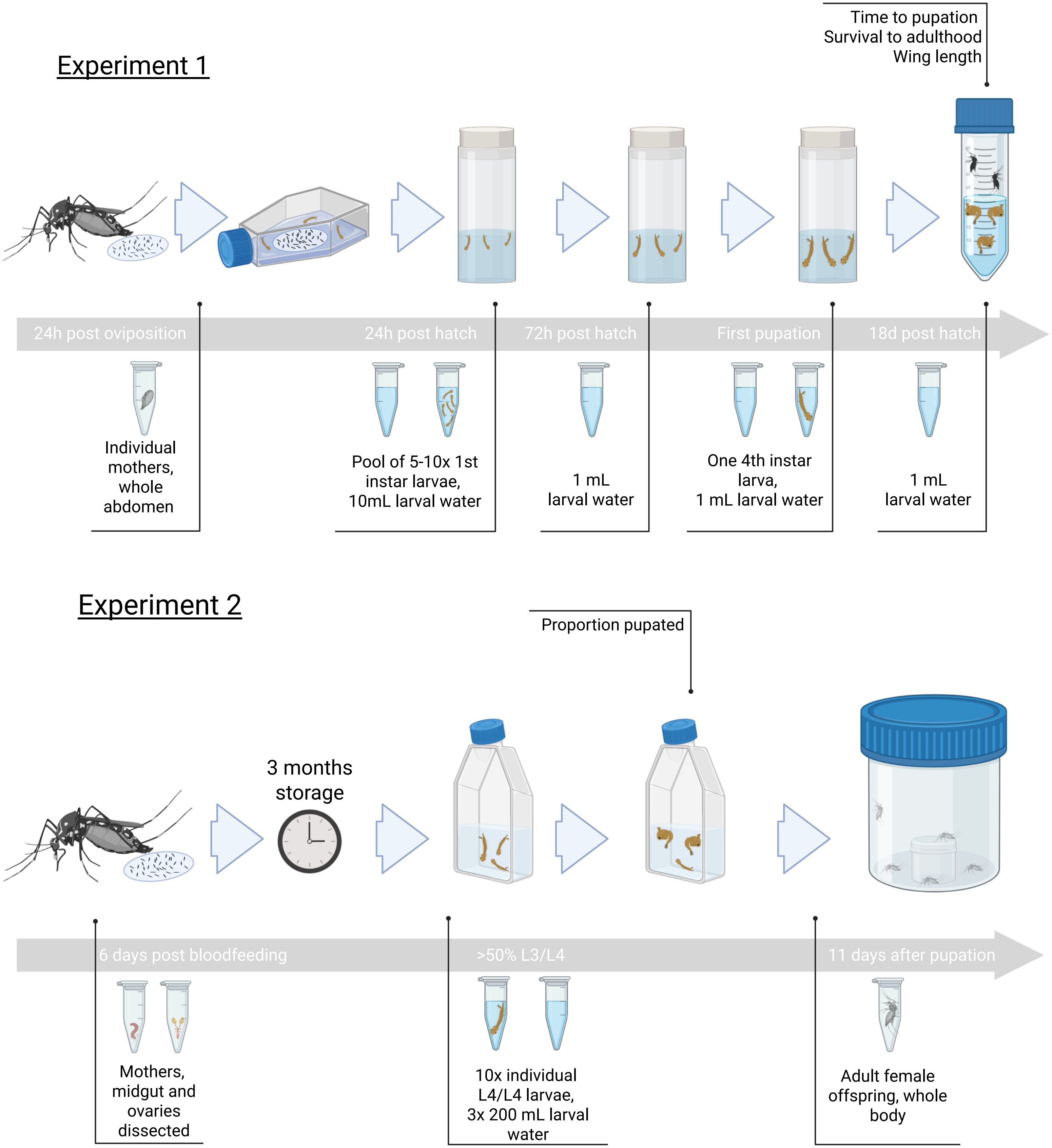

### af_2.pdf

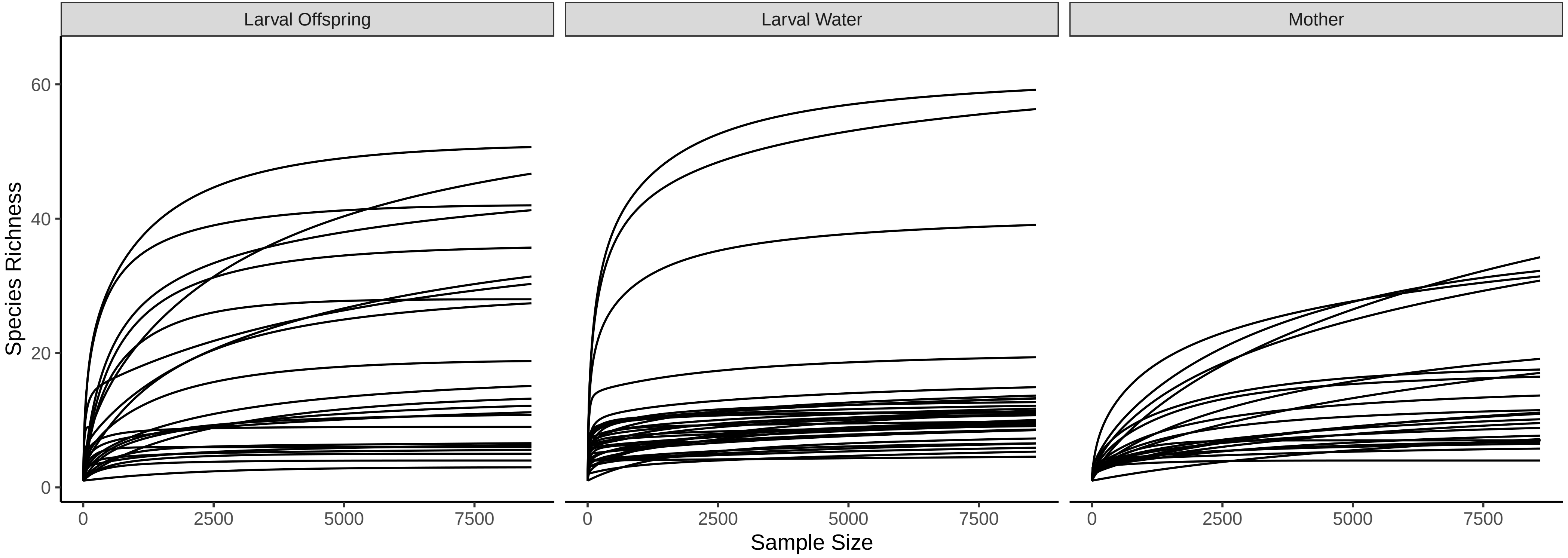

### af_3.pdf

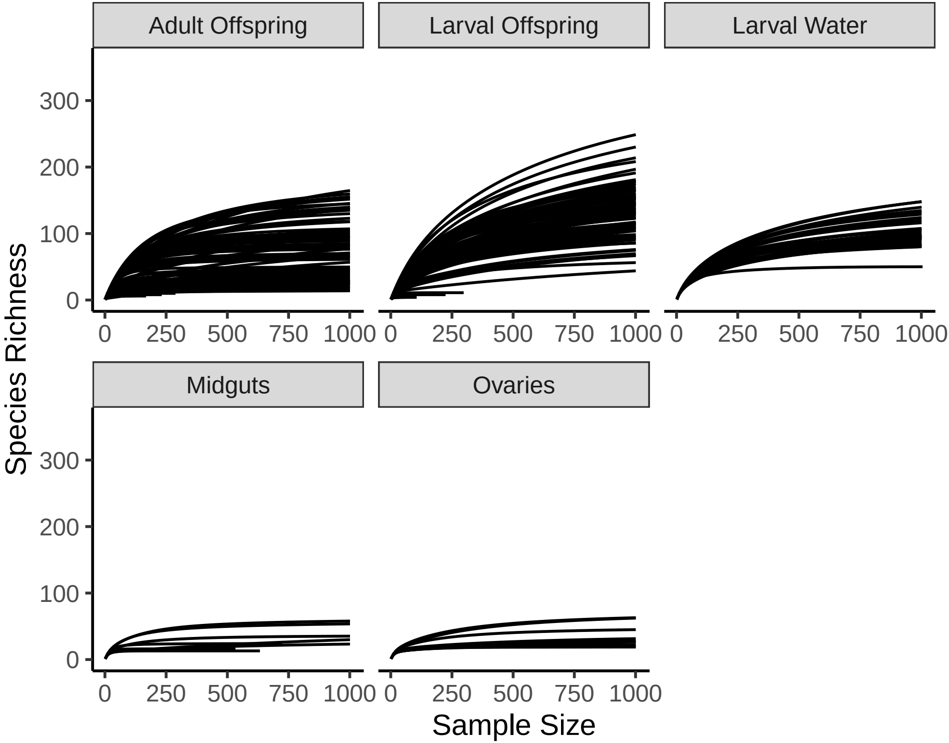

### af_4.jpg

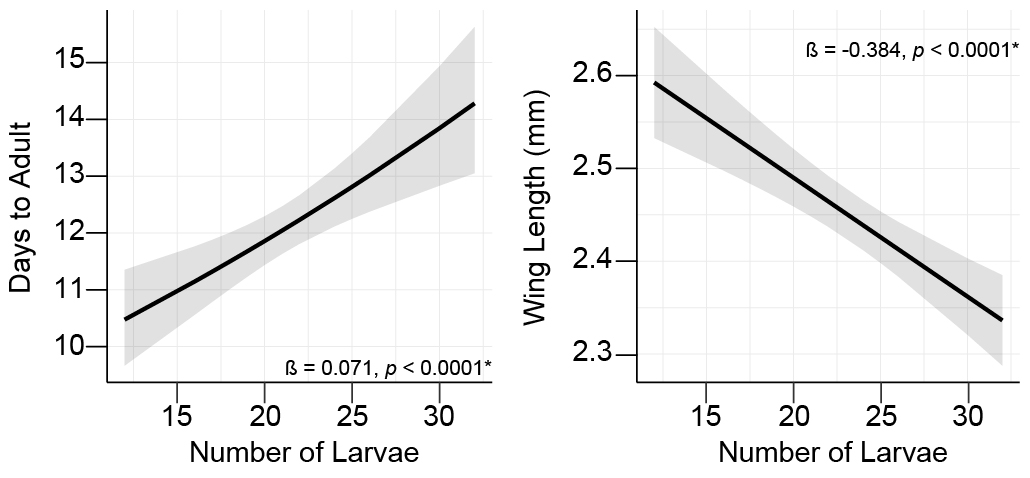

### af_6.pdf

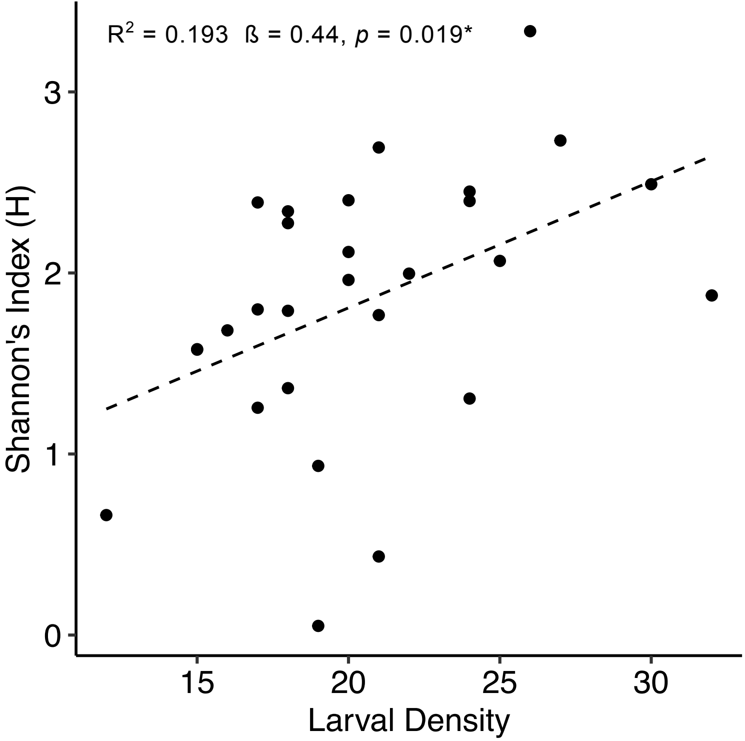

### af_8.pdf

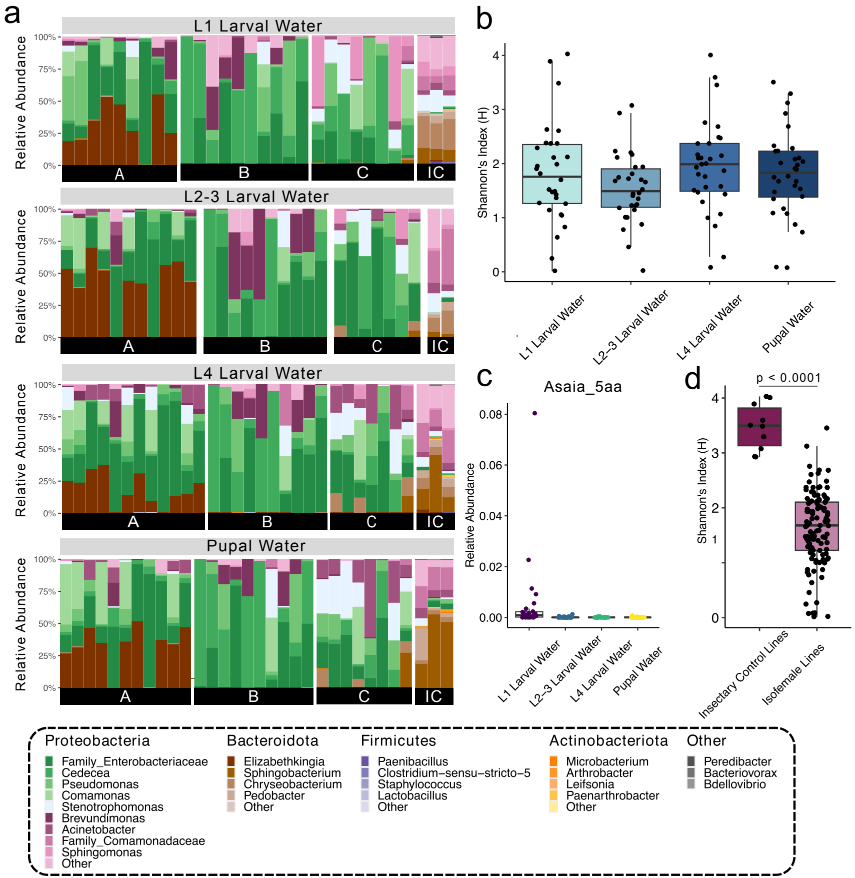

### af_10.png

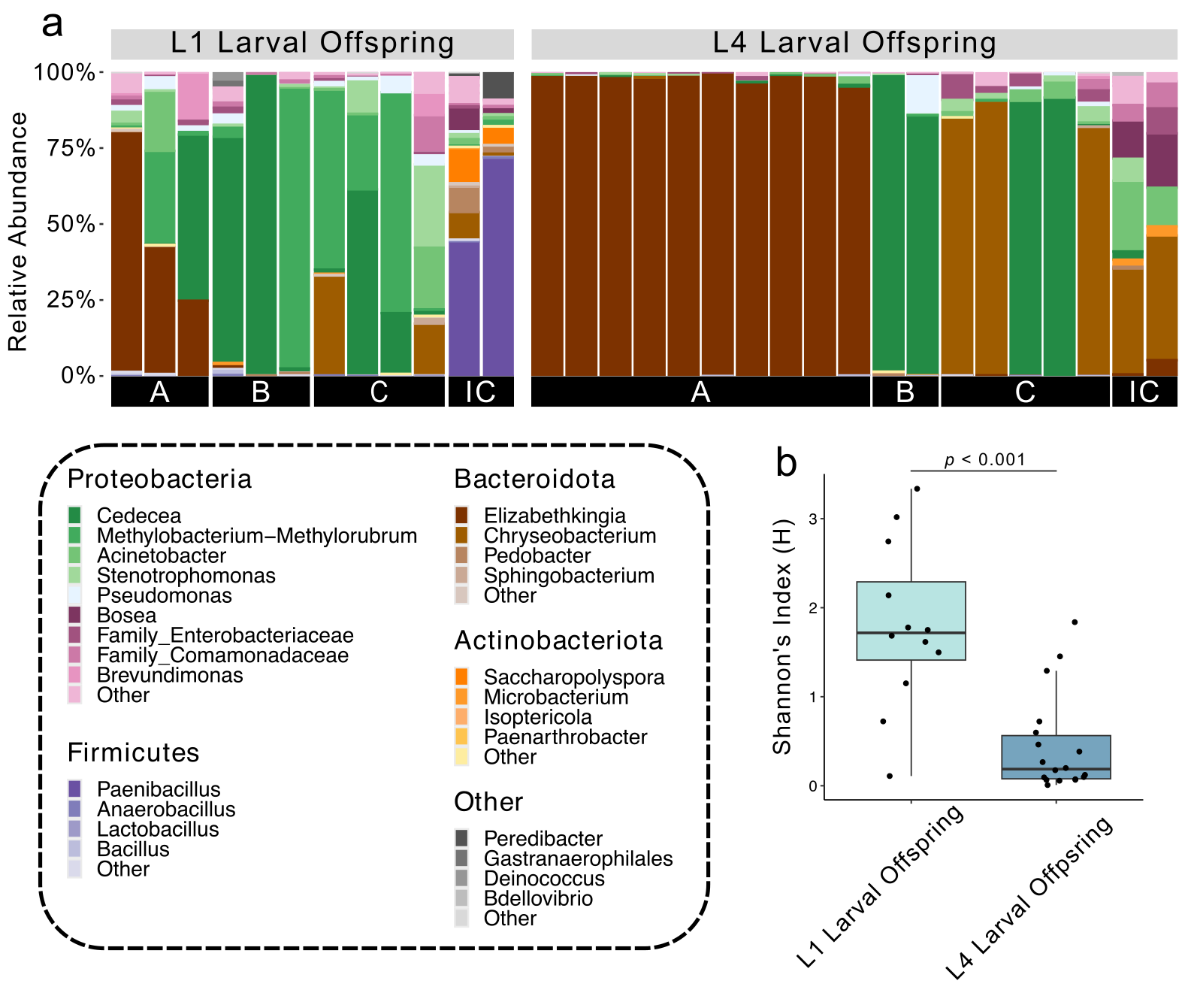

### af_12.pdf

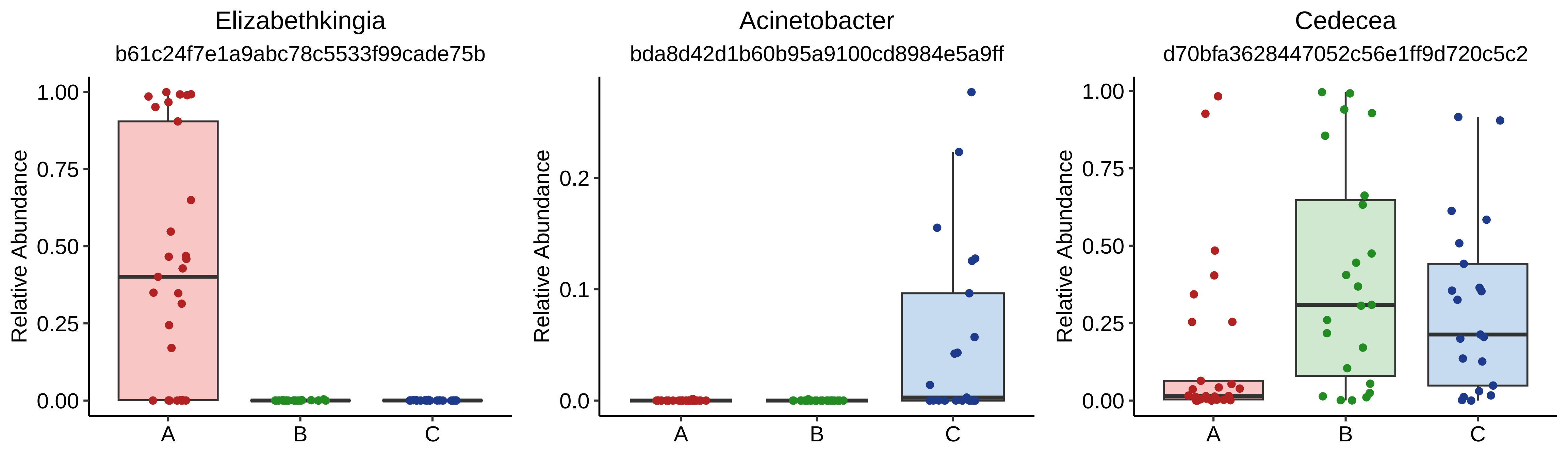

### af_14.pdf

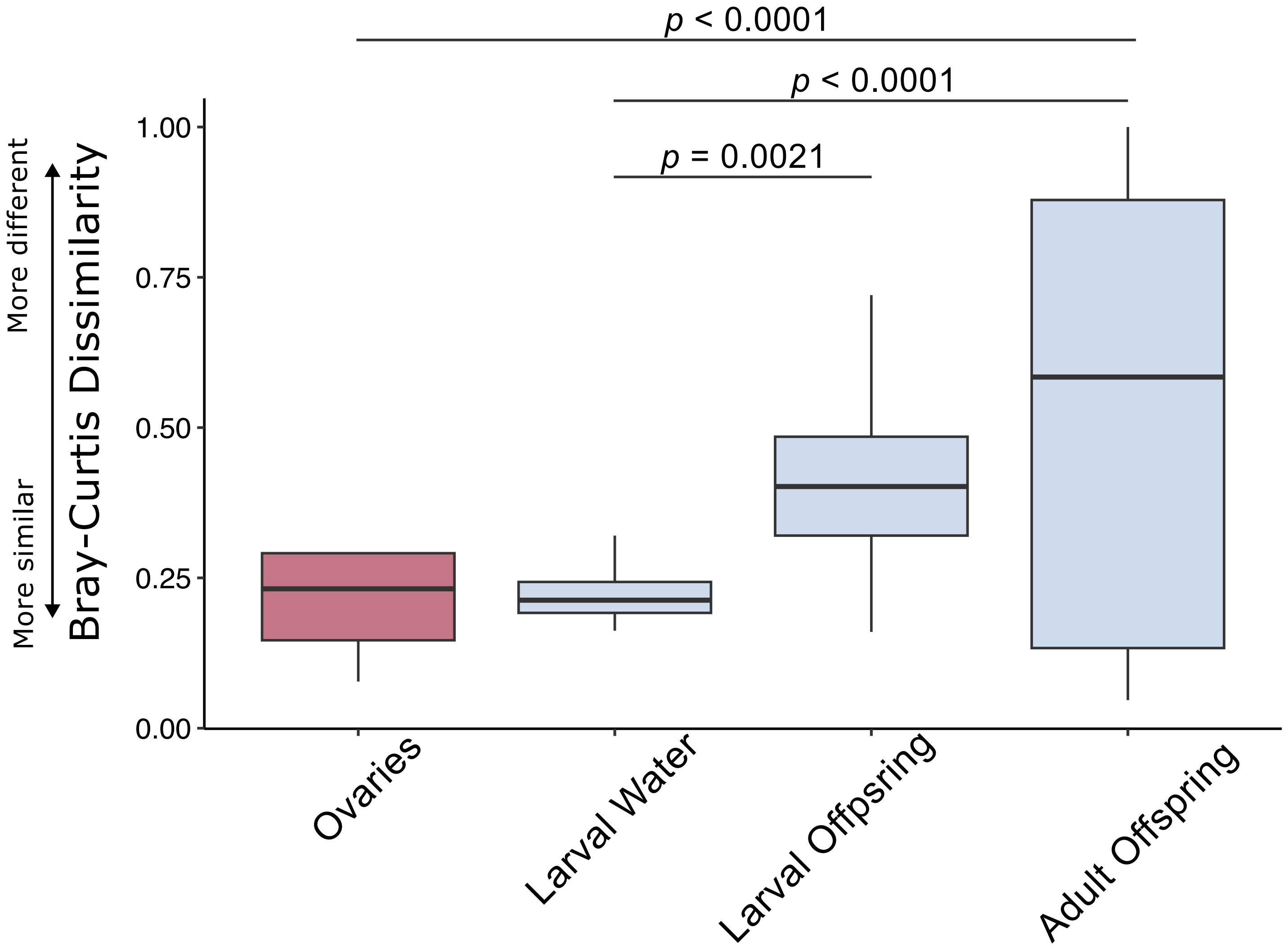

### af_15.pdf

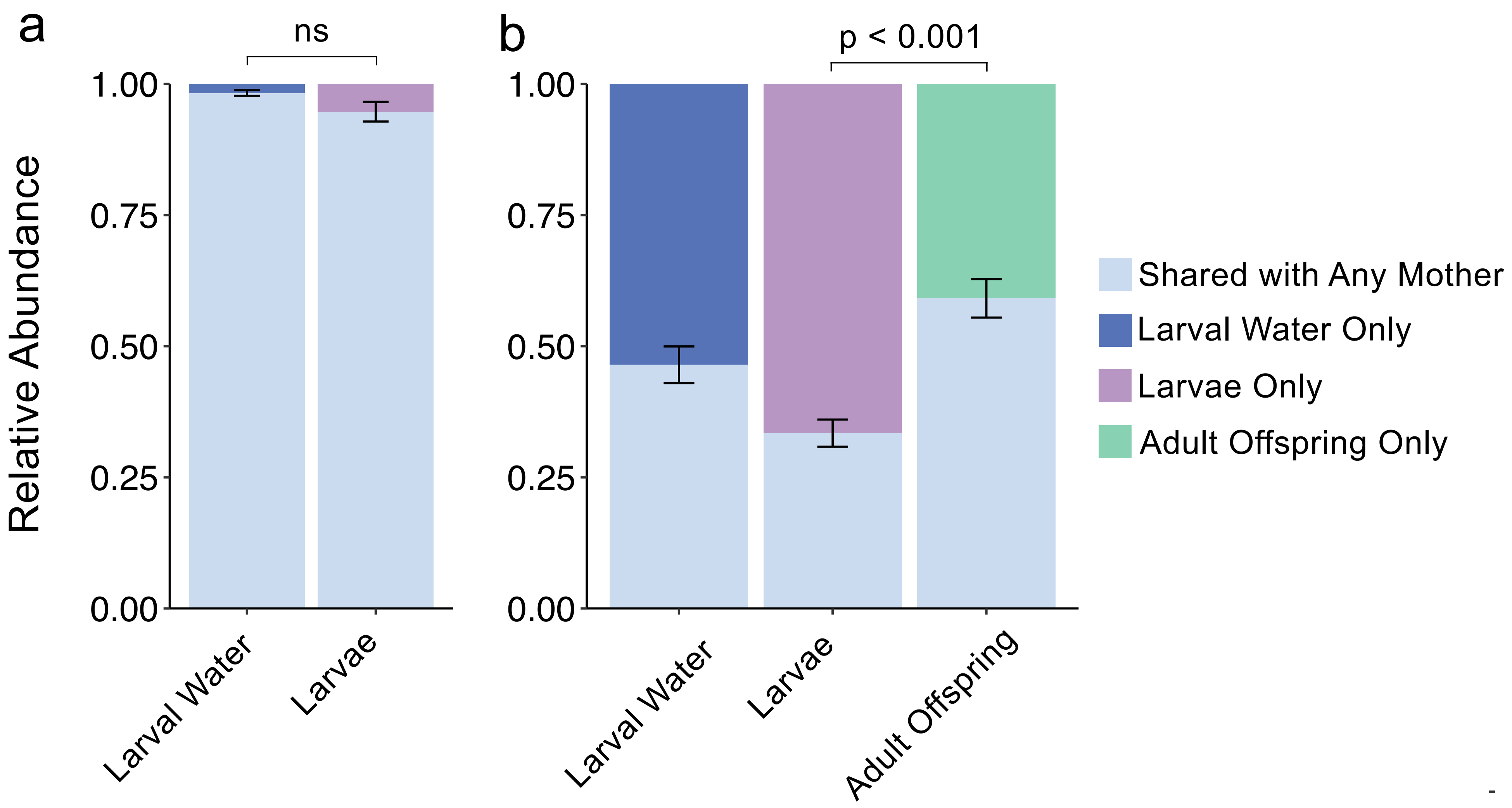

### af_16.pdf

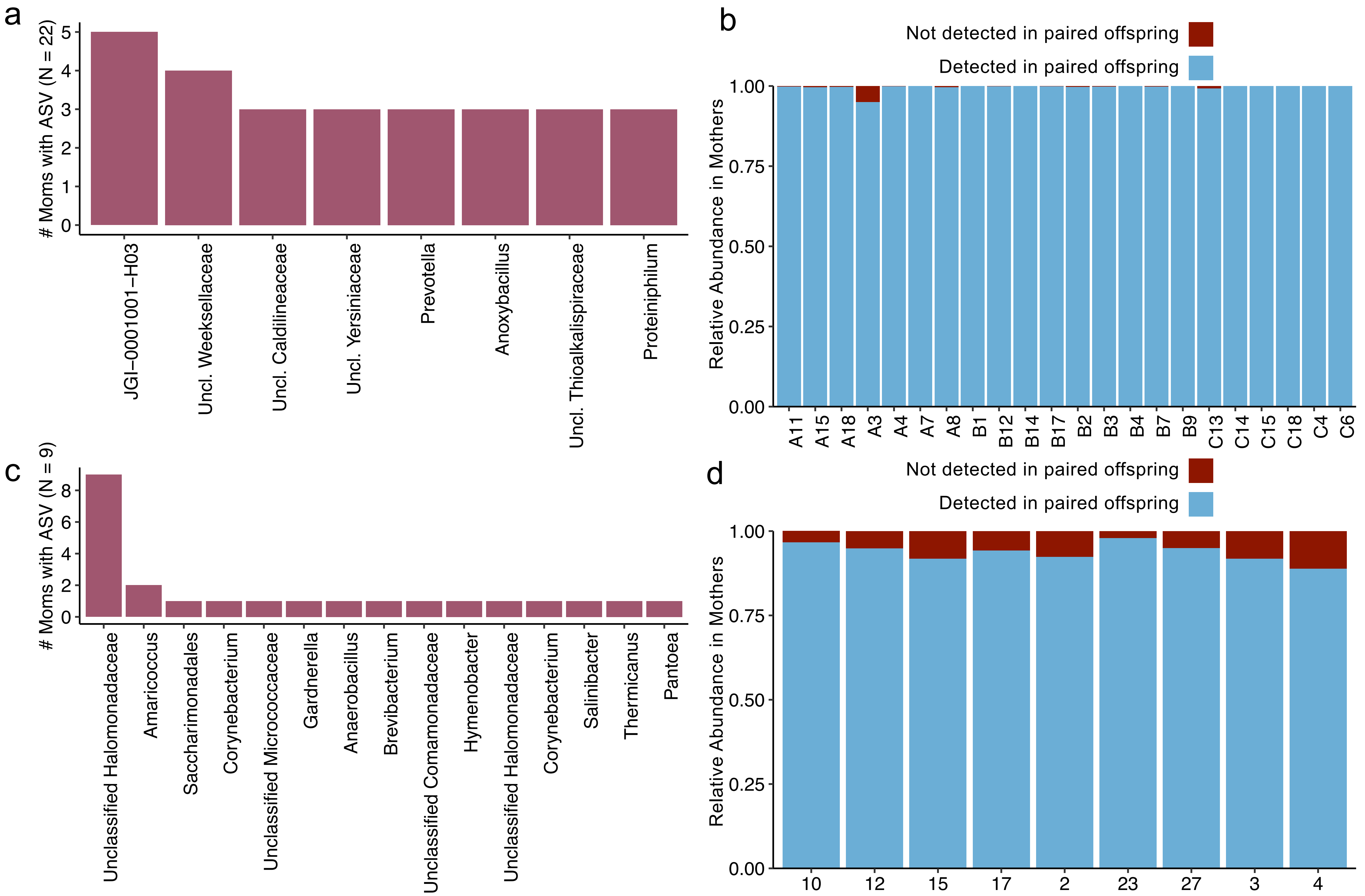

### af_17.pdf

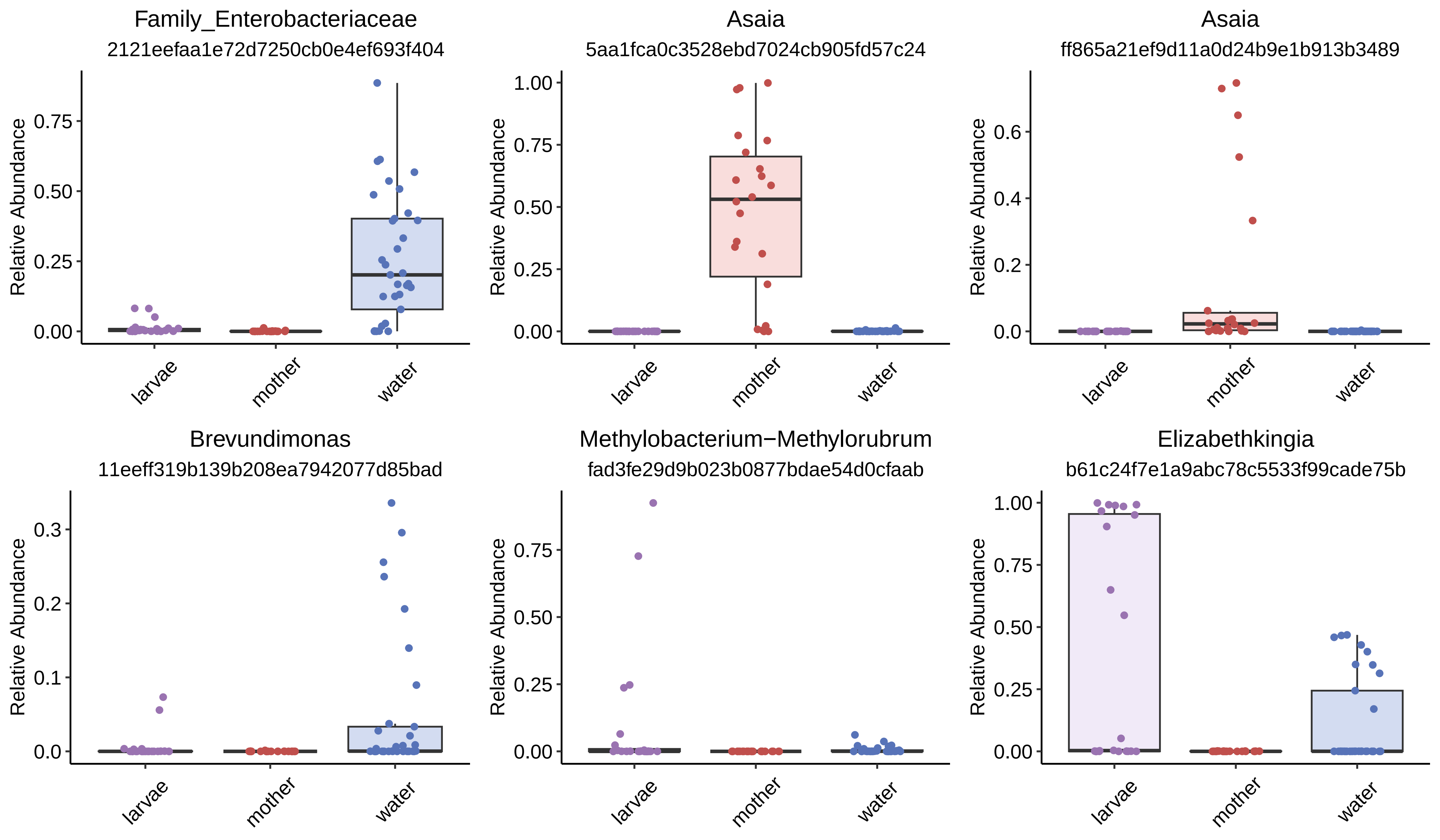

### af_18.pdf

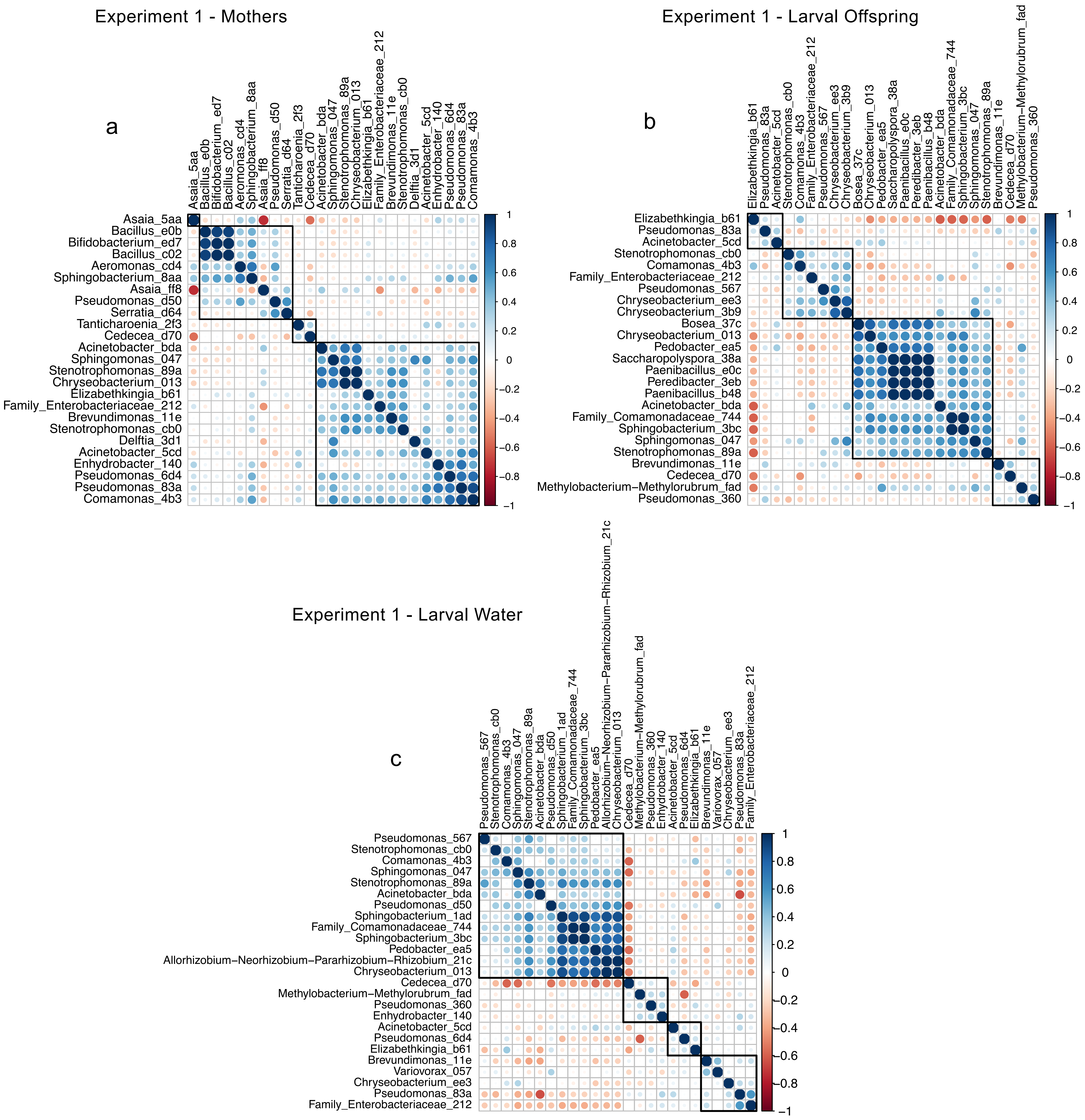

### af_19.pdf

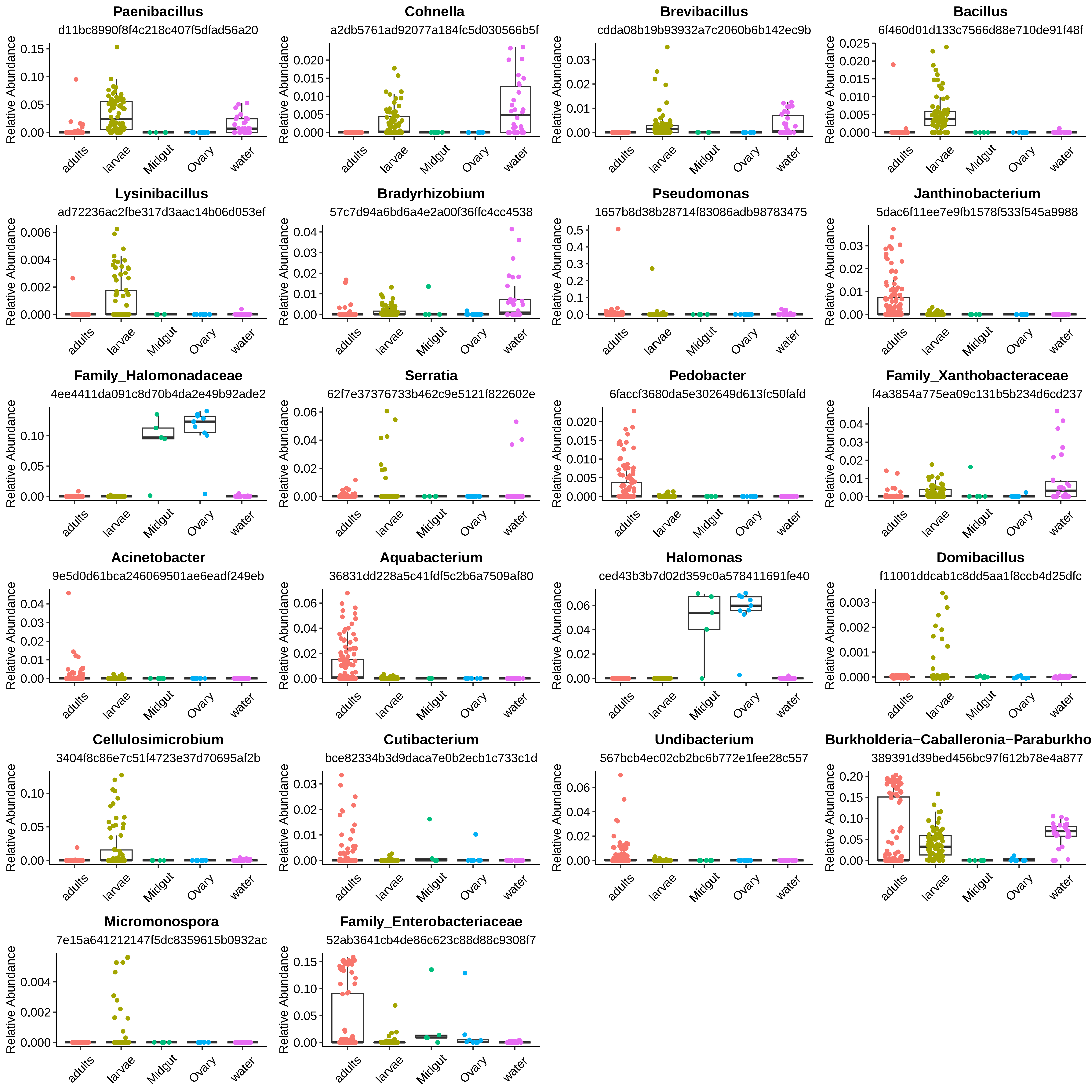

### af_20.pdf

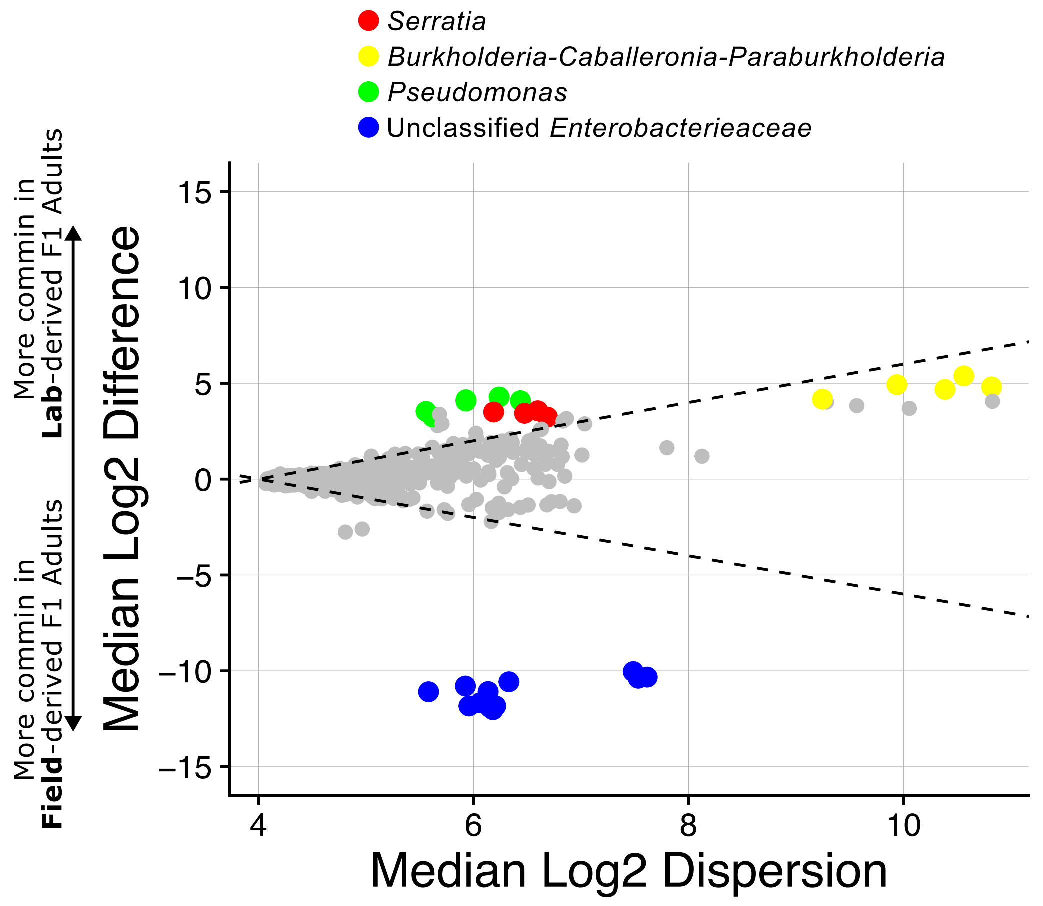

### af_21.pdf

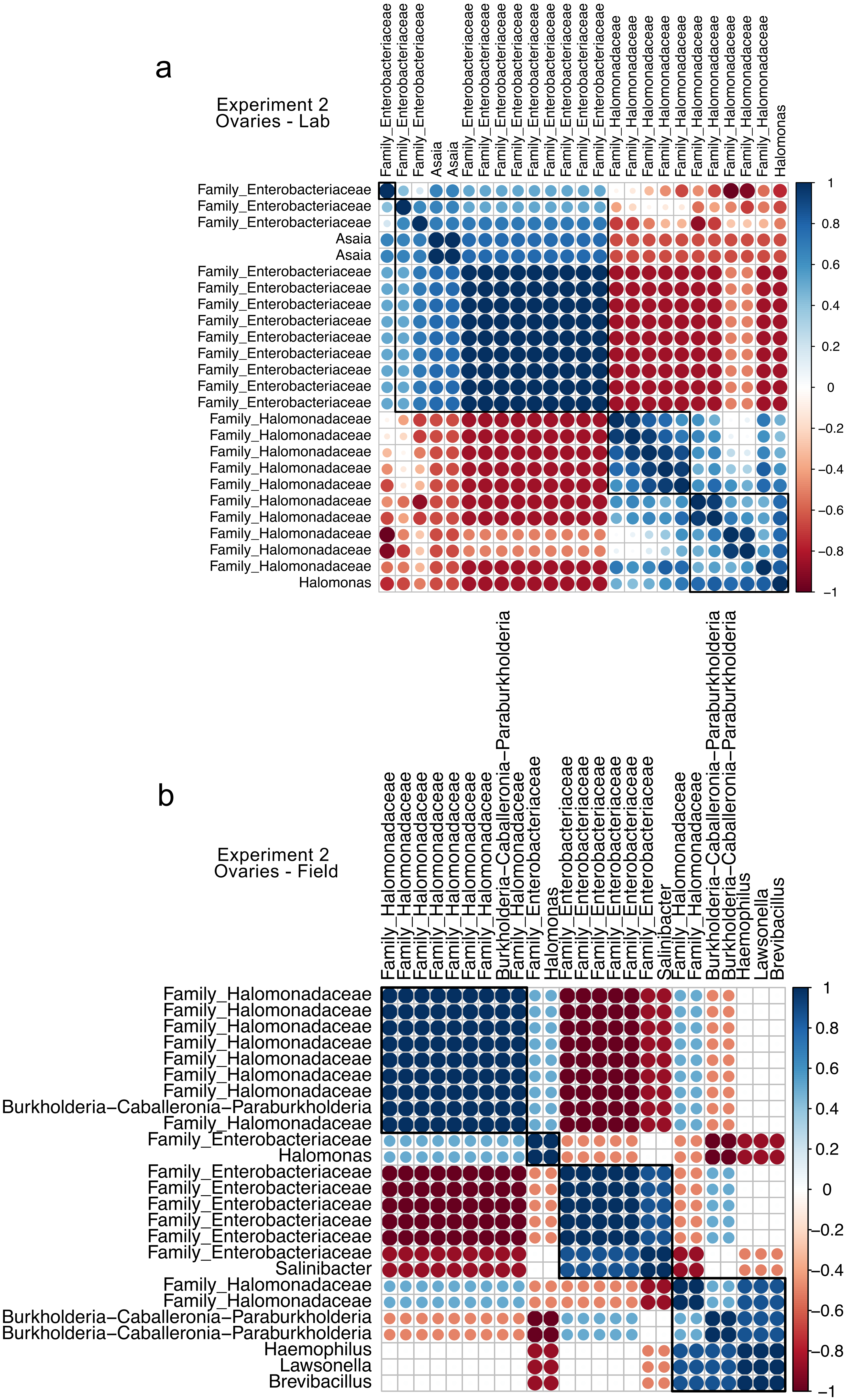

### af_22.pdf

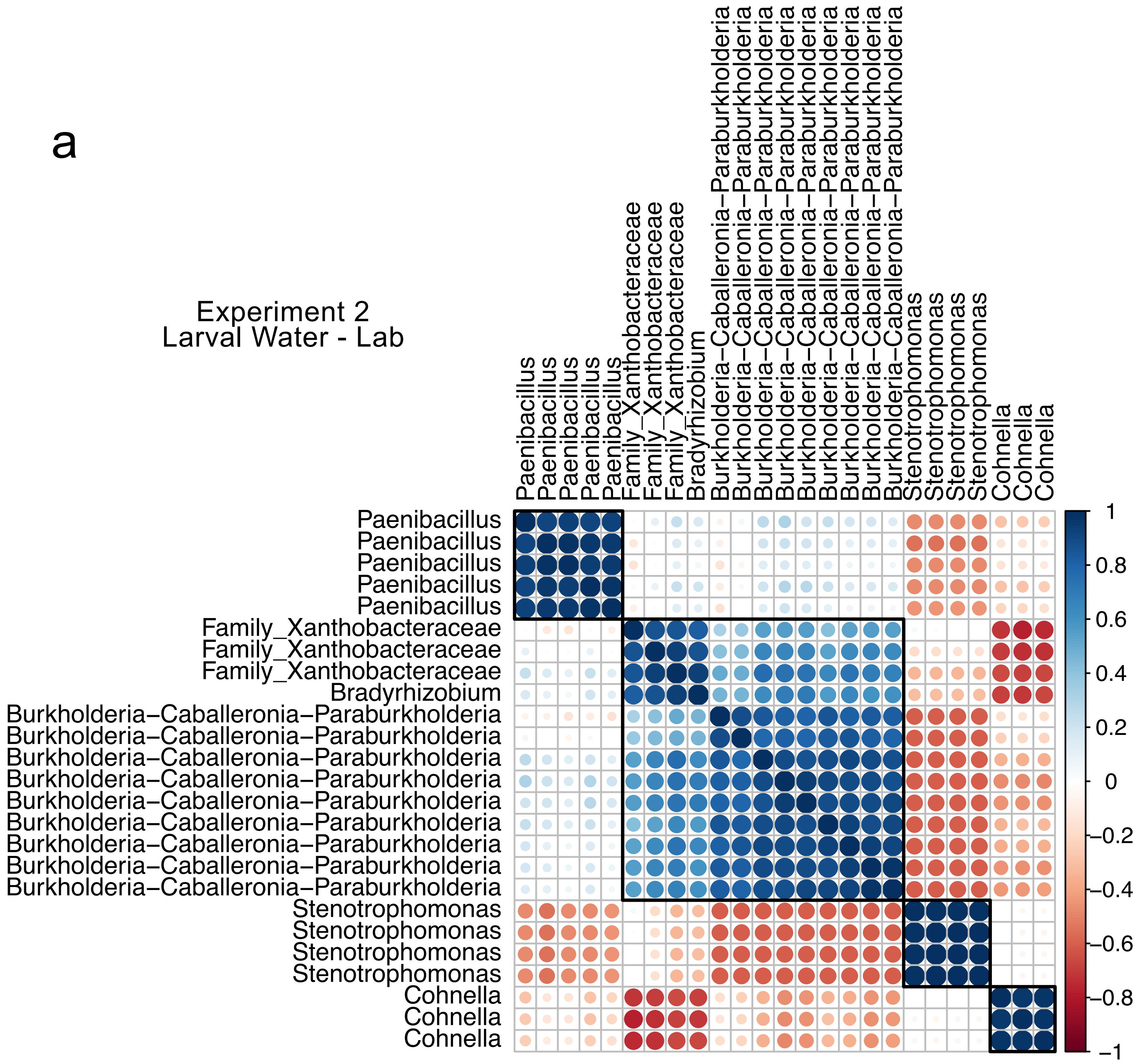

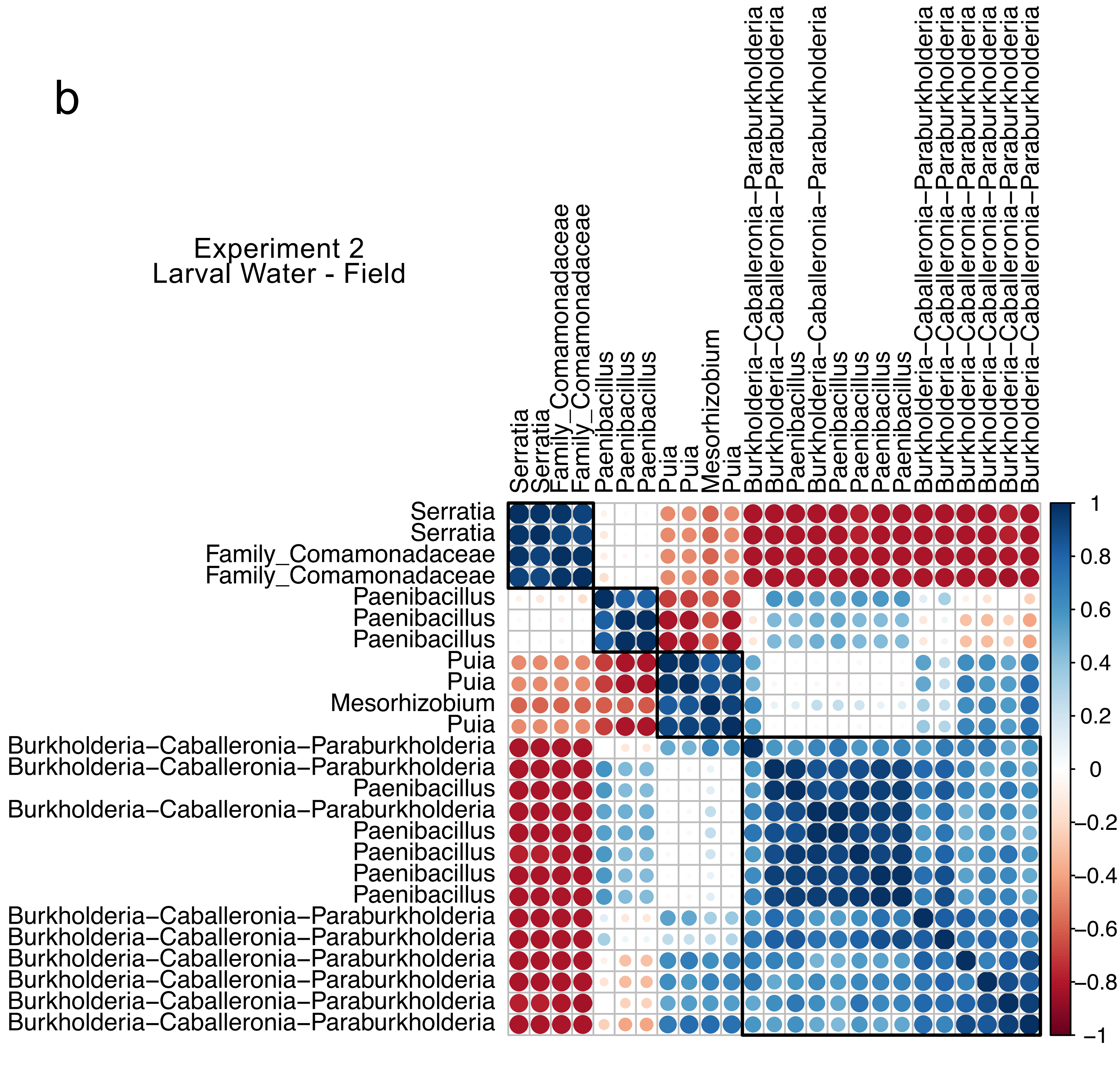

### af_23.pdf

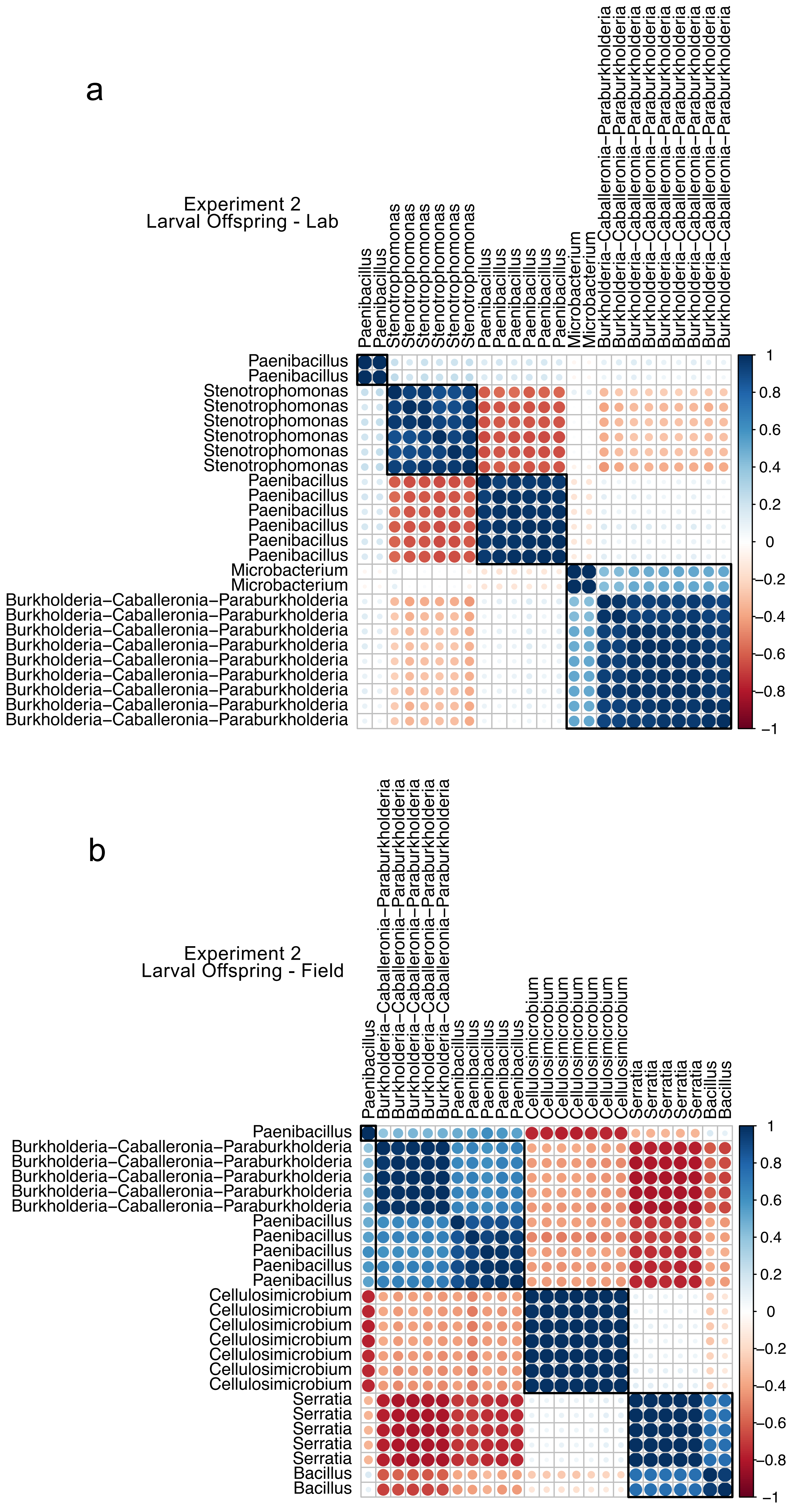

### af_24.pdf

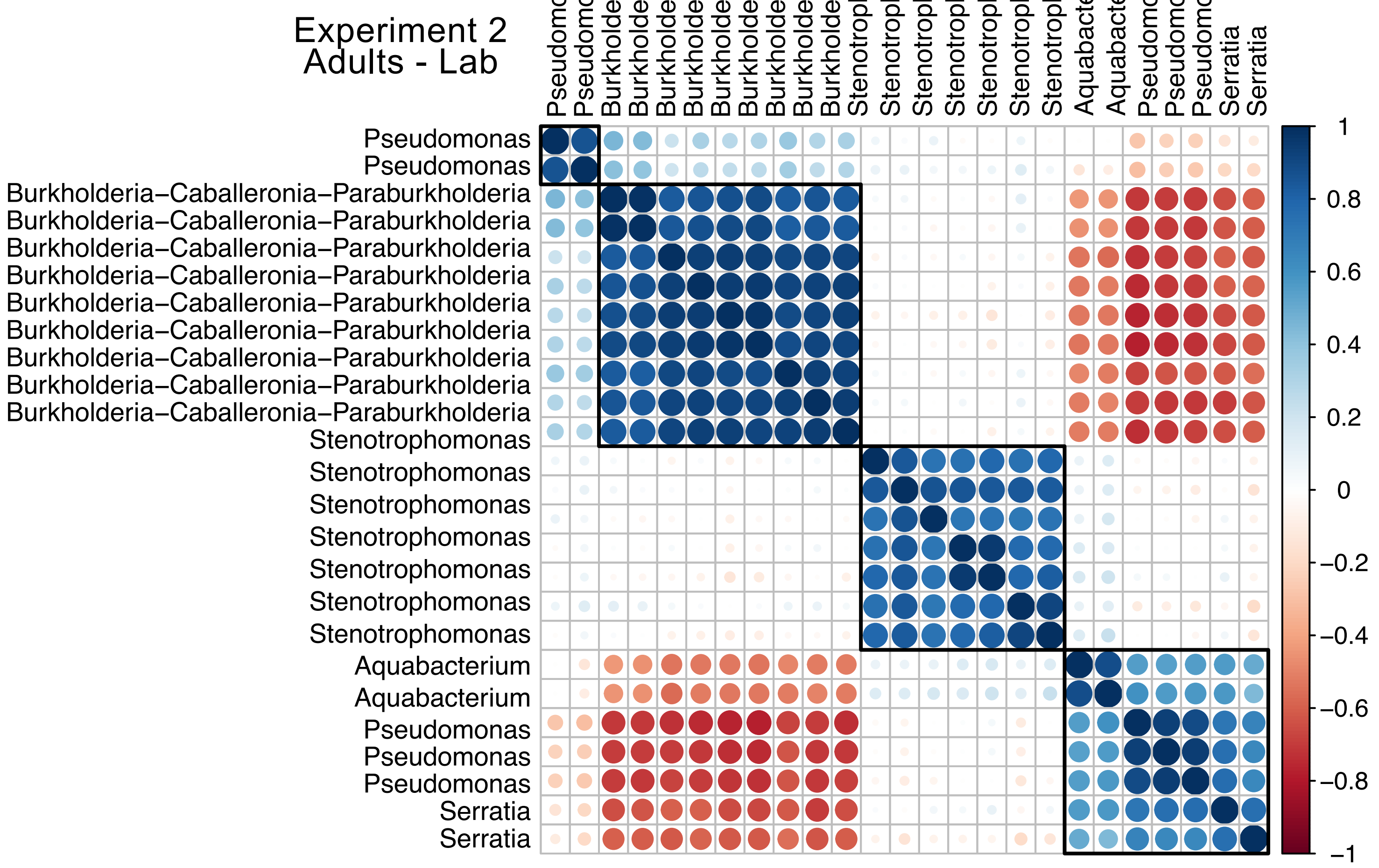

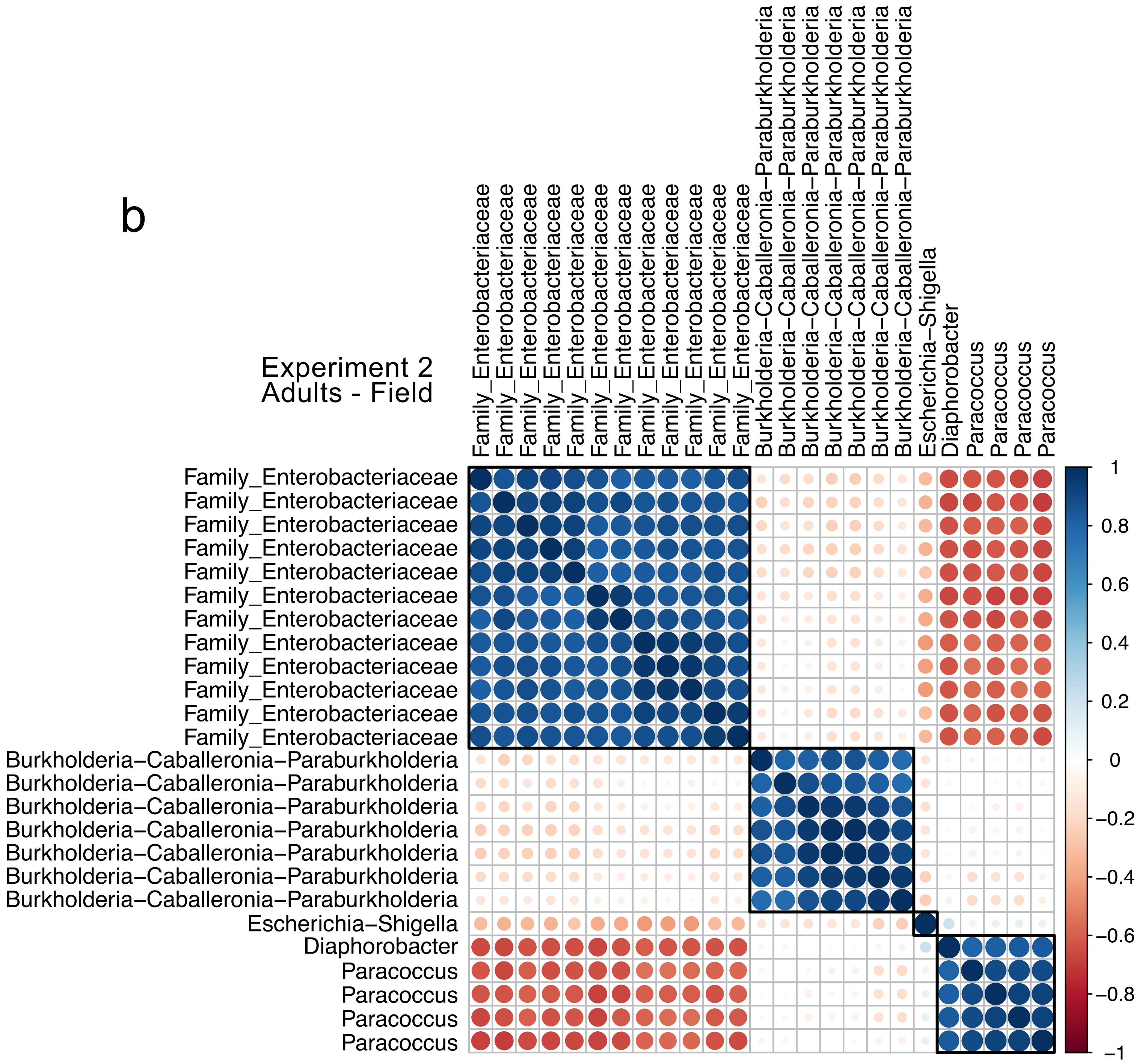

### af_25.pdf

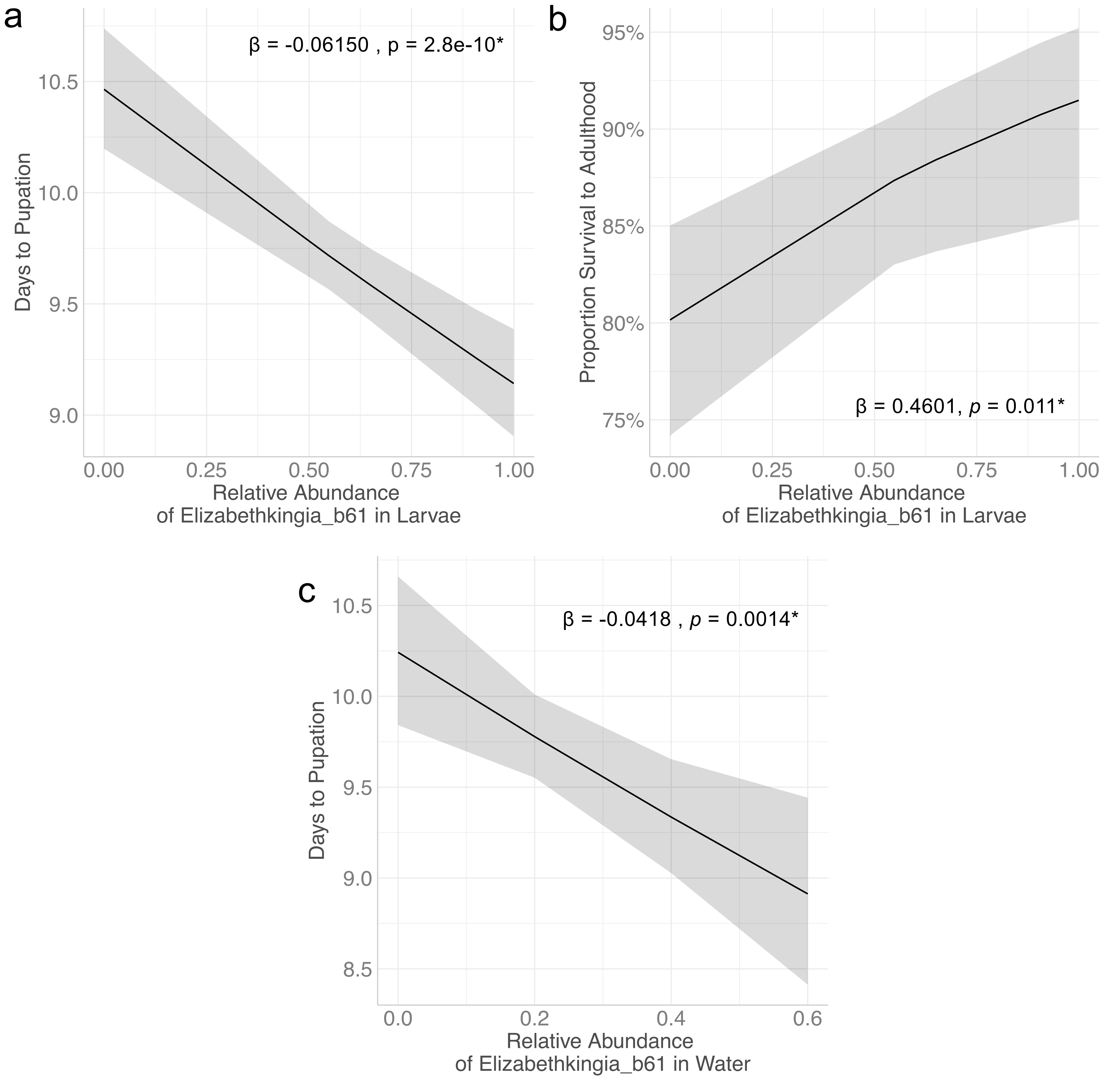

### af_26.pdf

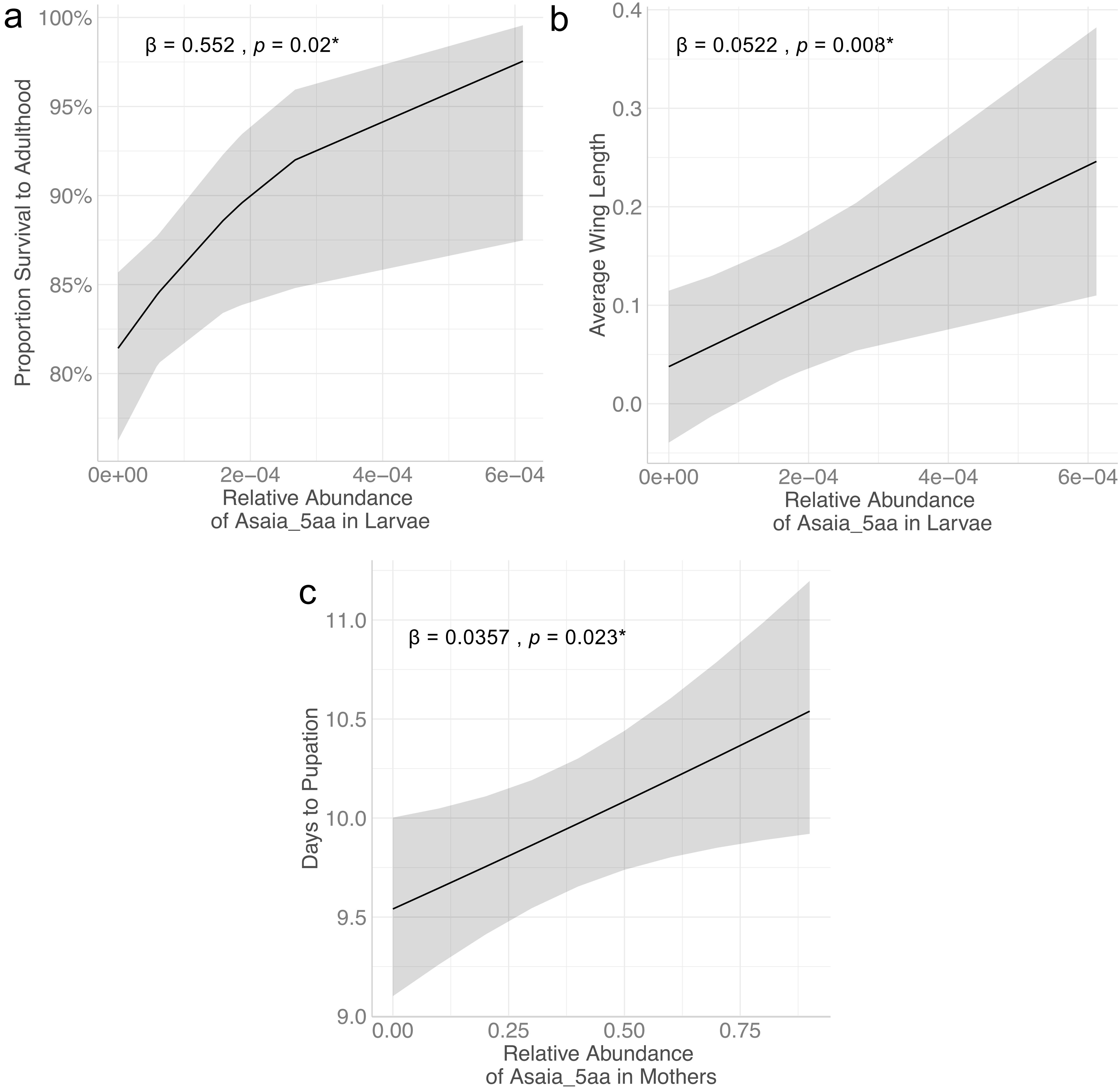

### af_27.pdf

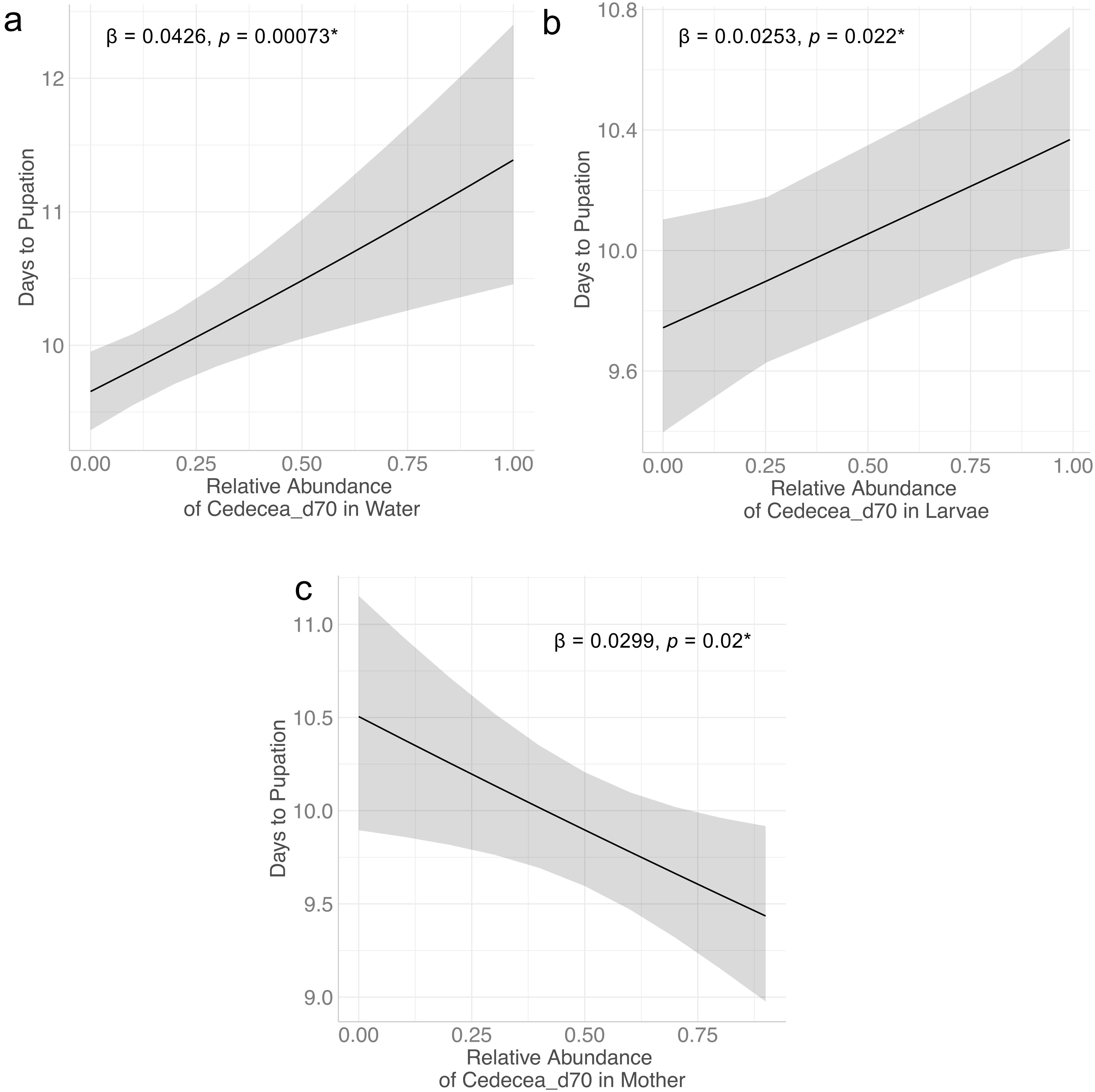

### af_29.png

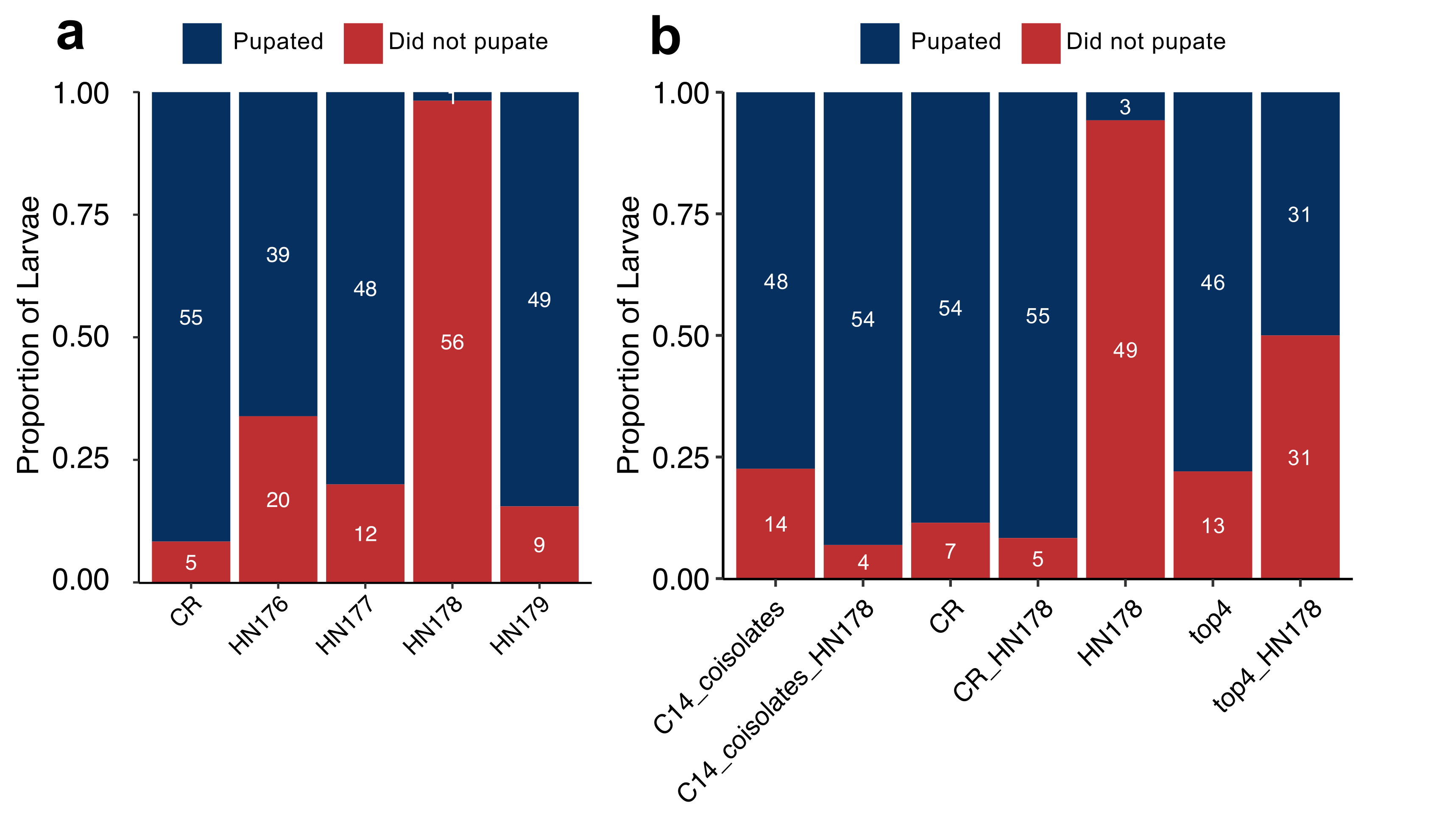

### af_31.pdf

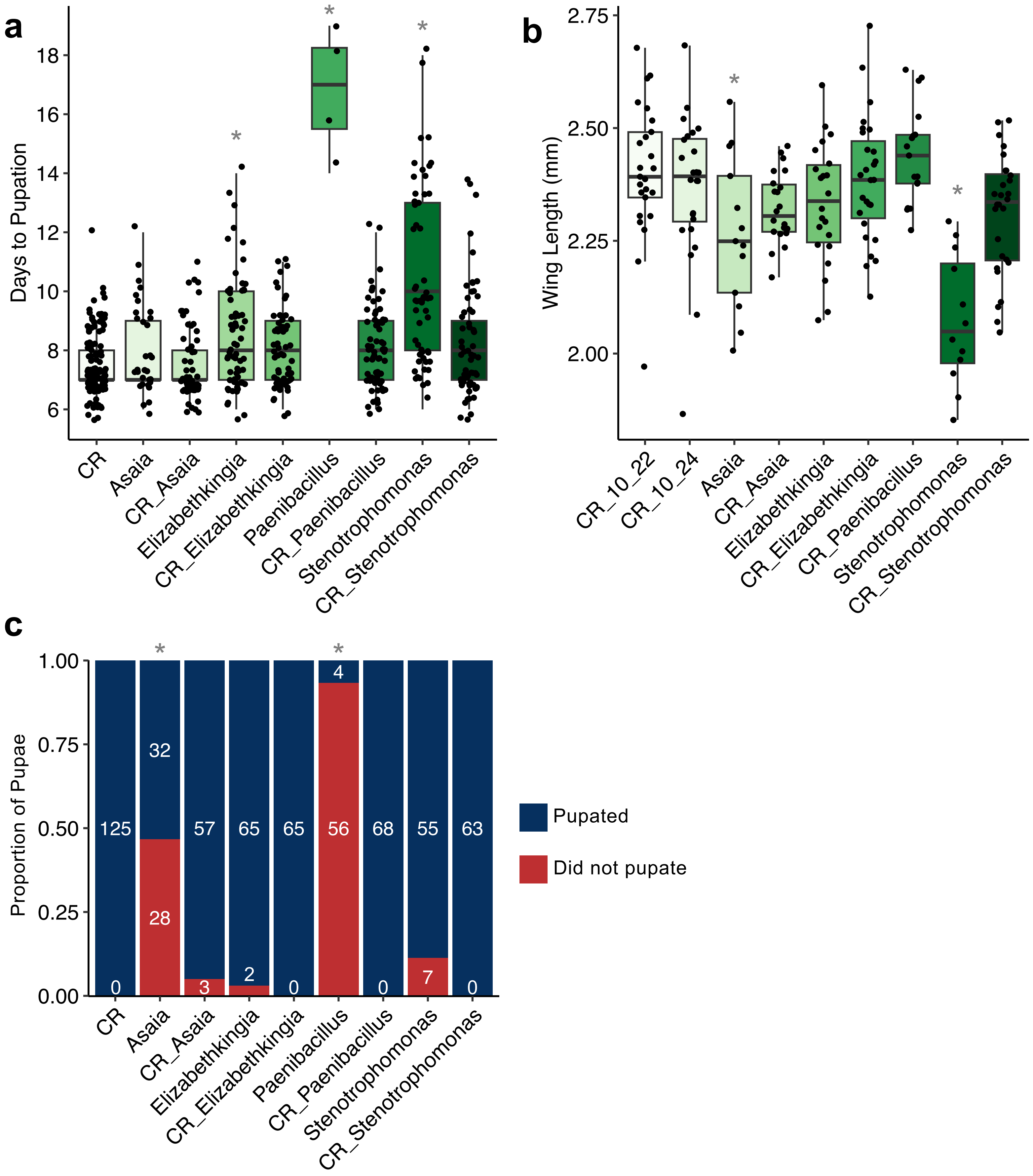
